# Structural basis for ring-opening fluorescence by the RhoBAST RNA aptamer

**DOI:** 10.1101/2024.12.30.630784

**Authors:** Shea H. Siwik, Aleksandra J. Wierzba, Shelby R. Lennon, Lukasz T. Olenginski, Amy E. Palmer, Robert T. Batey

## Abstract

Tagging RNAs with fluorogenic aptamers has enabled imaging of transcripts in living cells, thereby revealing novel aspects of RNA metabolism and dynamics. While a diverse set of fluorogenic aptamers has been developed, a new generation of aptamers are beginning to exploit the ring-opening of spirocyclic rhodamine dyes to achieve robust performance in live mammalian cells. These fluorophores have two chemical states: a colorless, cell-permeable spirocyclic state and a fluorescent zwitterionic state. Recently, the developed dye SpyRho555 almost exclusively adopts the closed state in solution and becomes fluorescent in complex with the RhoBAST aptamer. To understand the basis for RhoBAST-SpyRho555 fluorogenicity, we have determined crystal structures of RhoBAST in complex with 5-carboxytetramethylrhodamine and a SpyRho555 analogue, MaP555. RhoBAST is organized by a perfect four-way junction that positions two loops to form the dye-binding pocket. The core of the ligand resides between a tri-adenine floor and a single guanine base, largely driven by π-stacking interactions. Importantly, the unpaired guanine interacts with the 3-position group of MaP555 to stabilize the open conformation, supported by mutagenesis data, and may play an active role in promoting the open conformation of the dye.

**Graphical Abstract:** 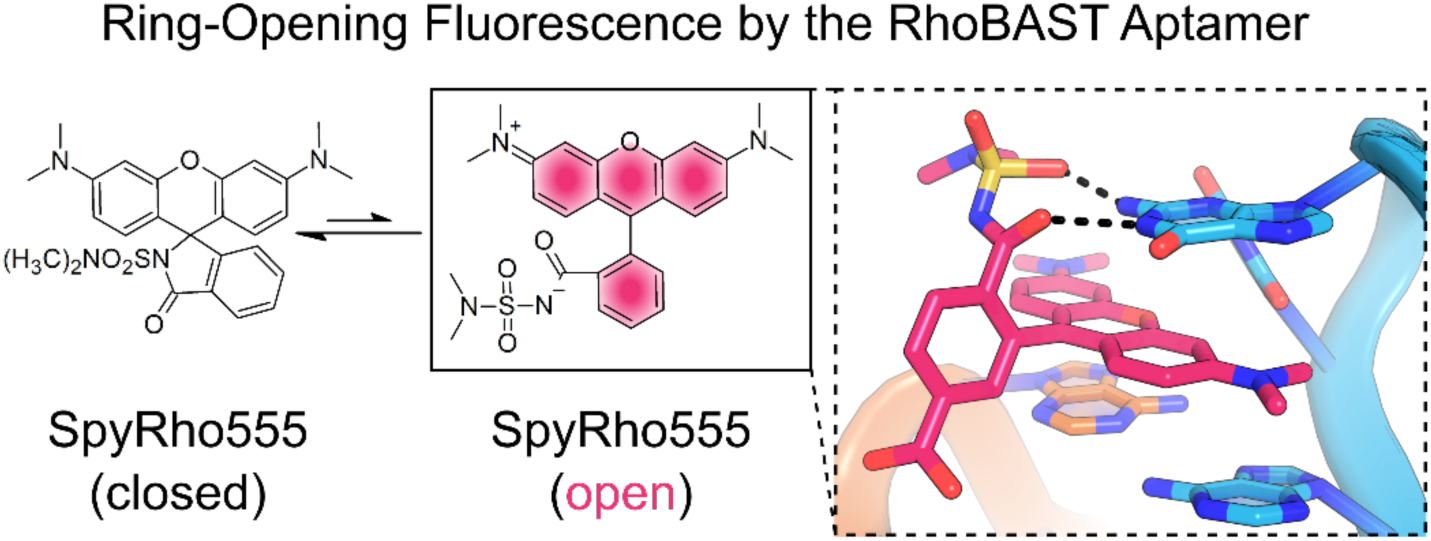

## INTRODUCTION

Imaging the transcriptome in live cells is essential to understanding RNA function by informing cellular localization and dynamics. Approaches that insert a tag within the RNA of interest (ROI), generally in the 3′ UTR, provide a fluorescent signal that can be visualized. Currently, the gold standard for RNA imaging is the MS2-MS2 coat protein (MS2-MCP) system (1, 2). This method typically inserts 24 copies of a small hairpin into the ROI that binds a dimer of the MCP with high affinity. MCP is expressed as a fluorescent protein (FP) fusion that accumulates fluorescent signal on the ROI allowing for single molecule visualization (3). Since its development, the MS2-MCP system has been used to gain diverse insights into RNA metabolism such as transcription of RNA in condensates (4), the effects of DNA supercoiling on transcription regulation (5), and the localization of lncRNA in neurons (6) by tagging ROIs. While this approach has been highly successful, considerations such as a high background signal due to unbound, diffusing FPs and large tags consisting of both RNA and protein constructs that alter endogenous RNA functions or localization (3, 7) need to be addressed. An alternative to the MS2-MCP system that could remedy these issues is the use of a fluorogenic RNA aptamer. These RNA elements have been *in vitro* selected to bind synthetic, small molecule fluorophores, thereby inducing fluorescence turn-on. Despite the potential for fluorogenic aptamers to be powerful tools to expand the capabilities of RNA imaging, these aptamers have failed to supplant the MS2-MCP system and achieve widespread use, indicating that continuing improvement and benchmarking of fluorogenic aptamer performance is needed.

The mechanism by which fluorogenic aptamers induce fluorescence is dictated by the nature of the fluorescent dye. Twisted intramolecular charge transfer (TICT) can occur in dyes that freely rotate in solution and fluoresce when rigidified in a planar configuration via π-stacking with RNA bases. Fluorogenic aptamers using this mechanism include Broccoli/Spinach (8, 9), Mango (10), and Peppers (11). A second mechanism is contact-quenching, which covalently links a fluorophore to a contact quencher to suppress fluorescence in solution and enhances signal when the two groups are physically separated upon RNA binding. Riboglow (12) and RhoBAST (13) are two examples of fluorogenic aptamers that use this mechanism. Both mechanisms have drawbacks when considering their implementation for widespread and robust RNA imaging. For example, fluorogenic aptamers that contain G-quadraplexes demonstrate poor folding in mammalian cells (14) and contact-quenching dyes lose dynamic range by never achieving the same level of fluorescence as the unconjugated fluorophore (12, 15).

An alternative mechanism of fluorogenicity that has gained traction in the imaging field involves exploiting spirolactonization or spirolactamization of rhodamines, a process we will refer to here as spirocyclization. Rhodamine dyes have long been valued for their good photophysical properties as demonstrated by widely used, commercial dyes such as rhodamine 123 and rhodamine 800 (16). The inherent propensity for rhodamines to spirocyclize establishes an equilibrium between two chemical states with only one being fluorescent (17, 18) (Figure 1A). The spirocyclic, closed form is cell-permeable and colorless while the zwitterionic, ring-open form is highly fluorescent. This equilibrium can be tuned by modifications to the amino groups of the xanthene ring (position 3′ and 6′), modification of the carboxyphenyl ring at positions 5 and 6, and modification of the carboxylic acid group (position 3) (19, 20). In addition, this equilibrium can be shifted by extrinsic factors such as solvent polarity, hydrogen bonding, metal ions, and pH (20–22). This property has been exploited in the development of next-generation fluorogenic tags where the rhodamine dyes are conjugated to HaloTag, SNAP-tag, or CLIP-tag and turn-on upon protein binding (23–25). Fluorogenic aptamers that can bind these rhodamine dyes and stabilize the ring-open form are attractive solutions to remedy some of the current problems associated with the other mechanisms primarily used by fluorogenic aptamers.

**Figure 1.**
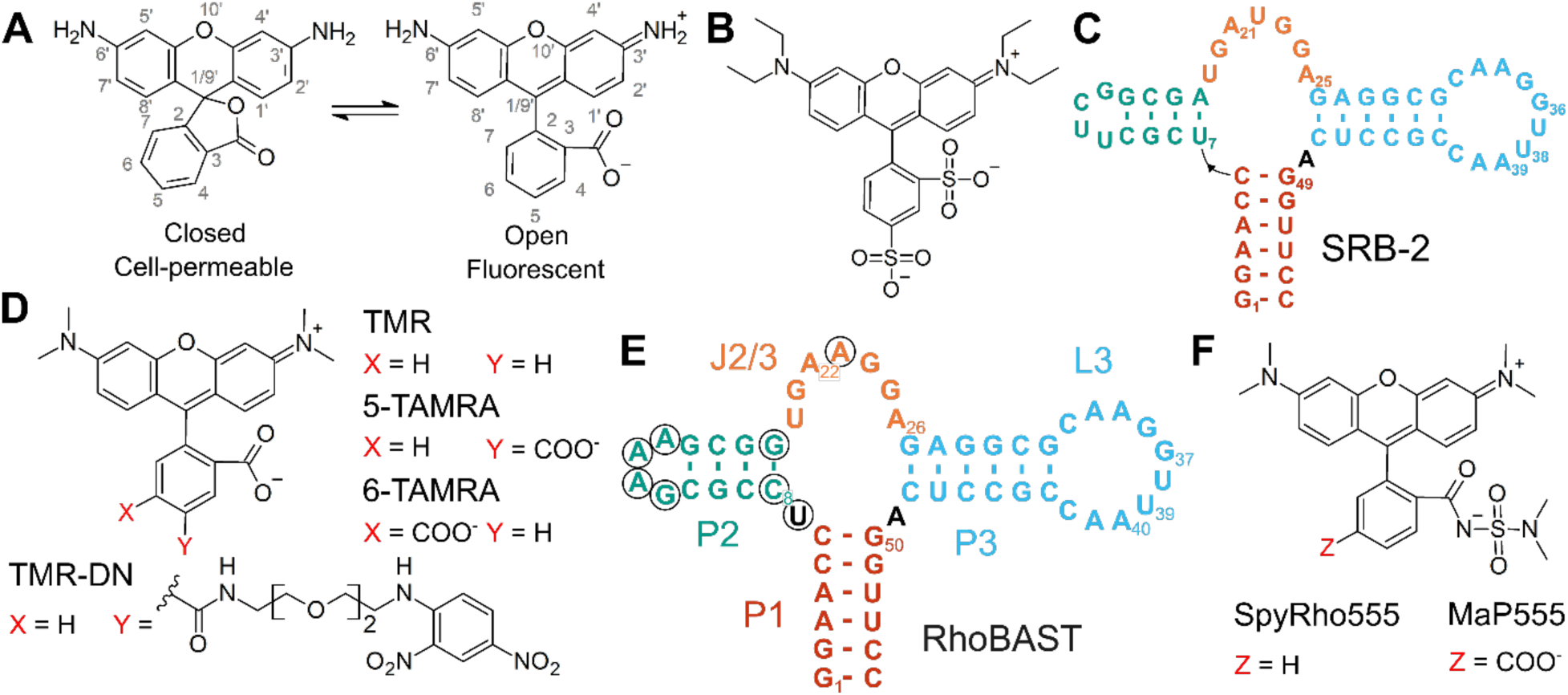
Chemical structures of relevant fluorophores and predicted secondary structures of the SRB-2 and RhoBAST aptamers. (**A**) Schematic demonstrating the spirocyclization equilibrium of rhodamine. The numbering scheme for rhodamines dyes is shown in gray. (**B**) Chemical structure of sulforhodamine B. (**C**) Predicted secondary structure of the SRB-2 aptamer. (**D**) Chemical structure of tetramethylrhodamine and related compounds. (**E**) Predicted secondary structure of the RhoBAST aptamer. Nucleotide positions that differ from the SRB-2 aptamer are circled. (**F**) Chemical structure of SpyRho555 and analogue MaP555.

The RhoBAST aptamer has emerged as an outstanding candidate for developing RNA tools that can bind spirocyclic rhodamines and increase fluorescence by ring-opening. RhoBAST derives from the SRB-2 aptamer originally selected to bind the fluorophore sulforhodamine B (26) (Figure 1B-C). This aptamer displayed the capability to bind analogous dyes with nanomolar affinity including tetramethylrhodamine (TMR) that, when conjugated to dinitroaniline (TMR-DN), can increase fluorescence using the contact-quenching mechanism (27) (Figure 1D). Because SRB-2 was not selected to bind TMR, a doped re-selection was performed against TMR to improve the aptamer’s performance, resulting in the RhoBAST aptamer (13). Overall, the sequences of SRB-2 and RhoBAST are highly similar with no changes occurring in the proposed binding regions (Figure 1E). Recently, the spirolactam rhodamine dye, SpyRho555, was developed for the RhoBAST aptamer, where RNA binding favors the open, fluorescent form of the dye (15). This dye is a derivative of TMR containing an *N, N*-dimethylsulfamide at the 3-position, identical to the amide group present in the fluorescent dye MaP555 (23) but lacking substitution at the 6-position of the phenyl ring (Figure 1F). Despite the bulkier 3-position group, RhoBAST binds the dye with high affinity (15). Future engineering of the RhoBAST aptamer:dye pair for diverse imaging applications will benefit from detailed structural insights, ensuring that essential structural elements and fluorogenic functionality are preserved.

To this end, we report the crystal structures of RhoBAST in complex with two rhodamine dyes: 5-carboxytetramethylrhodamine (5-TAMRA) and the SpyRho555 analogue, MaP555. The RNA structure reveals a perfect four-way H junction (4HJ) at the core of the RNA where helices P1 and P3 stack and J2/3 stacks on helix P2 in an antiparallel arrangement. Extensive tertiary interactions between J2/3 and P3 form the binding site, establishing a cleft at the terminal loops of these regions. The xanthene core of the dye π-stacks on an adenine shelf formed by a type II A-minor interaction, placing the 3-position carboxylate of 5-TAMRA for hydrogen bonding with a critical guanine, G31. Interactions between G31 and the 3-position are preserved in the co-crystal structure of RhoBAST with MaP555, indicating how RhoBAST stabilizes the zwitterionic conformation of SpyRho555. This work demonstrates how the RhoBAST aptamer engages with rhodamine dyes to stabilize the open configuration and provides structural information that can be used to engineer RhoBAST for imaging applications or in the creation of novel aptamers that increase fluorescence by ring-opening mechanisms.

## MATERIALS AND METHODS

### Materials

SpyRho555 was purchased from Cytoskeleton, Inc. (CY-SC018) or synthesized according to established protocols (15). The free acid form of MaP555 was purchased from Tenova Pharmaceuticals (T01218). 5-TAMRA and 6-TAMRA were purchased from Santa Cruz Biotechnology (sc-495643, sc-495644), and 5(6)-TAMRA was purchased from Novabiochem (8.51030).

### RNA Preparation

DNA oligonucleotides were commercially synthesized (Integrated DNA Technologies) and dsDNA templates for transcription were generated using recursive PCR (28). Sequences of all oligonucleotides used in this study are given in Supplementary Table S1. RNA was *in vitro* transcribed using T7 RNA polymerase as previously described (29). Transcribed RNA was subsequently purified using 10% denaturing PAGE (29:1 acrylamide:bis-acrylamide, 8 M urea, 1x TBE (90 mM Tris base, 90 mM boric acid, 3 mM ethylenediaminetetraacetic acid (EDTA)). Product RNA, as visualized by UV shadowing, was excised from the gel and eluted using the “crush-and-soak” method into 0.5x TE buffer (5 mM Tris pH 8.0, 0.5 mM EDTA) or Milli-Q H_2_O overnight at 4 °C. The supernatant was buffer exchanged into 0.5x TE or Milli-Q H_2_O and concentrated using a 3 kDa MWCO centrifugal concentrator (Amicon). The RNA was filtered using a 0.22 µm centrifugal filter (Corning) and stored at -20 °C until use. RNA concentration was determined by measuring the absorbance at 260 nm; RNA extinction coefficients were calculated using the NovoPro Oligo Calculation Tool.

### Crystallization and Structure Determination

RhoBAST:5-TAMRA crystals were grown at 20 °C using the hanging drop vapor diffusion method by combining 1 µL of RNA solution (310 µM RhoBAST RNA and 900 µM 5(6)-TAMRA) with 2 µL of mother liquor (40 mM sodium cacodylate pH 7.0, 12 mM sodium chloride, 80 mM potassium chloride, 5% v/v 2-methyl-2, 4-pentanediol (MPD), and 20 mM spermine tetrahydrochloride) over 500 µL of 35% v/v MPD. After two days of growth, 3 µL of soaking solution (40 mM sodium cacodylate pH 7.0, 12 mM sodium chloride, 80 mM potassium chloride, 20 mM spermine tetrahydrochloride, 10% v/v MPD, 2 mM iridium(III) hexammine chloride, and 900 µM 5(6)-TAMRA) was added to the drop and incubated for 30 minutes before flash freezing looped crystals in liquid nitrogen. Data were collected at the Advanced Light Source at Lawrence Berkeley National Laboratory using beamline 8.2.2 at 100 K (1.09720 Å). Two datasets collected from this crystal were scaled and merged with XDS (30). Phases and initial map were determined using AutoSol (31) in PHENIX (32). The model was built in Coot (33) and underwent iterative rounds of refinement using PHENIX (34). RNA local geometry was further optimized using ERRASER (35) prior to a final round of refinement. The final R_work_ and R_free_ were 0.216 and 0.241, respectively. Final collection and model statistics are given in Supplementary Table S2.

RhoBAST:MaP555 crystals were grown at 20 °C using hanging vapor drop diffusion. 1 µL of the RNA solution (300 µM RhoBAST aptamer RNA and 1200 µM MaP555) was combined with 2 µL of mother liquor (40 mM MES pH 6.0, 10 mM barium chloride, 80 mM potassium chloride, 25% v/v MPD, and 12 mM spermine tetrahydrochloride) over 500 µL of 35% v/v MPD. Crystals were harvested after five days and flash frozen in liquid nitrogen. Data were collected at the Macromolecular X-ray Crystallography Core at the University of Colorado Boulder with a RIGAKU XtaLAB MM003 and Dectris PILATUS 200K 2D hybrid pixel array detector at 100 K (1.54178 Å). Datasets were integrated and scaled using HKL 3000 (36). Molecular replacement was performed with Phaser (37) using the RhoBAST:5-TAMRA structure (PDB ID: 9BUN) as the starting model. Model refinement and processing remained the same as previously described except for performing refinement to account for twinning using the twin operator, h, -k, -h-l. The final R_work_ and R_free_ values were 0.254 and 0.271, respectively. Final collection and model statistics are given in Supplementary Table S2. Binding surface areas of the ligands were calculated using PDBePISA (38).

### Fluorescent Binding Measurements

Reactions were conducted at room temperature in 1x Aptamer Selection Buffer (13) (20 mM HEPES pH 7.4, 125 mM KCl, 1 mM MgCl_2_) with 0.01% v/v Tween-20 using constant concentrations of dye (5 nM for SpyRho555, 10 nM for MaP555, 20 nM for 5-TAMRA, 20 nM for 6-TAMRA). The RNA was snap-cooled by heating to 75 °C for two minutes followed by cooling to 25 °C at a rate of -0.1 °C/second prior to use as described previously for this aptamer (13). Reactions were equilibrated for 1 hour in the dark at room temperature before pipetting reactions into a 384-well Low Flange microplate (Corning) and taking measurements using a CLARIOstar plus microplate reader (BMG Labtech). A reaction containing no dye or RNA was used for background subtraction. Measurements were taken using the following excitation and emission wavelengths and bandwidths: SpyRho555 (552-8 nm/585-12 nm), MaP555 (524-16 nm/584-8 nm), and 5-TAMRA/6-TAMRA (477-10 nm/575-20 nm). Data were fit to the quadratic equation

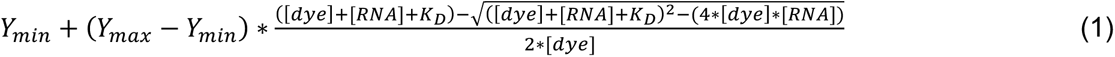

where Y_min_ is the lower baseline, Y_max_ is the upper baseline, [dye] is the concentration of fluorophore used for the experiment, [RNA] is the concentration of RNA, and K_D_ is the dissociation constant. Data were plotted and fitted using KaleidaGraph (v5.04).

### Chemical Footprinting

RhoBAST RNA was chemically probed using selective 2′-hydroxyl acylation analyzed by primer extension (SHAPE) with *N*-methylisatoic anhydride (NMIA) as the chemical probe (39). Probing reactions were carried out on RNA folded as previously described (40). Specifically, RNA was heated to 75 °C for 2 minutes and cooled to 60 °C at a rate of -0.1 °C/s. At 60 °C, 5x Aptamer Selection Buffer (100 mM HEPES pH 7.4, 625 mM potassium chloride, 5 mM magnesium chloride) and 5(6)-TAMRA dissolved in dimethyl sulfoxide (DMSO), SpyRho555 dissolved in DMSO, or neat DMSO were added to the RNA. The mixture was then cooled to 30 °C at a rate of -0.1 °C/second. Once at 30 °C, NMIA dissolved in DMSO or neat DMSO were added to the mixtures, and the reaction proceeded at 37 °C for 45 minutes. The final 10 μL probing reactions had concentrations of 100 nM RNA, 20 mM HEPES pH 7.4, 125 mM potassium chloride, 1 mM magnesium chloride, 10 µM ligand, and 13 mM NMIA. Reactions were reverse transcribed without a prior ethanol precipitation step, as previously described (41) using a ^32^P-labeled reverse transcription primer. Products were run on a 12% denaturing polyacrylamide gel at 55 W for 3 hours. The gel was exposed to a storage phosphor screen and imaged using an Amersham Typhoon 5 (Cytiva), and the image was analyzed using the SAFA software package (42).

### Absorption and Emission Spectra of Dyes

All data were collected in 1x Aptamer Selection Buffer. The spectra of the free dye were measured for MaP555, 5-TAMRA and 6-TAMRA at a concentration of 1 µM. The same dye concentrations were used to collect data in the presence of 5 µM RhoBAST. The RNA was heated at 90 °C for 3 minutes and then incubated on ice for at least 10 minutes prior to mixing with the dye. The RNA:dye mixture was incubated in the dark for 30 minutes before data collection. Absorption spectra were collected using a Cary 60 UV-Vis spectrophotometer. Absorption values were background corrected by subtracting the absorption values of a buffer control at each wavelength. The corrected and normalized absorbance values were plotted against wavelength in GraphPad Prism. Extinction coefficients (ε) were calculated based on the absorption values at λ_max_.

Fluorescence emission spectra were collected using QM-6 Steady-State Fluorimeter (PTI). All species were excited at the isosbestic point determined from their respective absorption spectra and are as follows: MaP555 +/- RNA (excitation 562 nm, emission range 565-620 nm), 5-TAMRA +/- RNA (excitation 556.5 nm, emission range 560-620 nm), 6-TAMRA without RNA (excitation 561 nm, emission range 564-620 nm), and 6-TAMRA with RNA (excitation 561 nm, emission range 570-620 nm). Fluorescence values were background corrected by subtracting the fluorescence values of a buffer control at each wavelength. The corrected and normalized fluorescence values were plotted against wavelength in GraphPad Prism.

### Docking Analysis of Closed Form of SpyRho555 with RhoBAST

The SpyRho555 ligand was manually built from the ligand in the RhoBAST:MaP555 co-crystal structure (PDB ID: 9DXL) in PyMol (v3.0.3, Schrödinger) in its ring open and closed states. The ligand coordinates were saved as an SDF file and imported into Molecular Operating Environment (MOE, v2022.02, Chemical Computing Group). Chain A of the RhoBAST aptamer domain (PDB ID: 9BUN) was used as the RNA receptor and was prepared for docking with the default Quick Prep protocol in MOE. Rigid receptor docking was carried out in a manner where the ligand search space was restricted to 4.5 Å surrounding the ligand in the RhoBAST:5-TAMRA co-crystal structure (PDB ID: 9BUN). The ligand was allowed to search 30 poses but only the top five were scored. Binding surface areas of the ligands were calculated using PDBePISA (38).

## RESULTS

### Crystallization of a RhoBAST:5-TAMRA complex

The predicted secondary structure of the RhoBAST aptamer, originating from the SRB-2 aptamer selected to bind sulforhodamine B, centers around a three-way junction motif. The highest degree of nucleotide conservation observed during selections of this aptamer resides within J2/3 and terminal loop L3 while helices P1 and P2 and terminal loop L2 vary in both length and composition. Using this information, we modified the RhoBAST aptamer to promote crystallization in the following ways. First, the lengths of the P1 and P2 helices were minimized to four and two base pairs, respectively, as we have observed in previous RNA:ligand complex crystallization efforts that smaller RNAs tend to be more favorable for crystallization (43). Second, we added an unpaired adenosine at the 3′-end which often promotes intermolecular lattice contacts. To validate that these changes were not deleterious to RNA function, we determined the equilibrium dissociation constant (K_D_) for SpyRho555 binding to the variant used for crystallization. We obtained a K_D_ of 70 ± 20 nM for both the wild type (WT) and RhoBAST variant, indicating that these changes do not affect the RNA’s ability to bind ligand (Table 1, Supplemental Figure S1A).

**Table 1.** RhoBAST K_D_ values.

| RNA Construct | Dye Identity | $K_D \pm \text{s.d. (nM)}$ | $K_{\text{rel}}^{\text{a}}$ |
| --- | --- | --- | --- |
| WT | SpyRho555 | $70 \pm 20$ | - |
| WT | 5-TAMRA | $82 \pm 4$ | 1.2 |
| WT | 6-TAMRA | $130 \pm 20$ | 1.9 |
| WT | MaP555 | $120 \pm 20$ | 1.7 |
| P1-4/P2-2 | SpyRho555 | $70 \pm 20$ | 1.0 |
| G31A | SpyRho555 | $> 30,000$ | $> 430$ |
| G31C | SpyRho555 | $> 45,000$ | $> 640$ |
| G19C | SpyRho555 | N.D.<sup>b</sup> | N/A |
| G19C/C27G | SpyRho555 | $\geq 2,300 \pm 500$ | 33 |
| U14C/A20G | SpyRho555 | $60 \pm 20$ | 0.86 |
| U14G/A20C | SpyRho555 | $180 \pm 30$ | 2.6 |
| C4U | SpyRho555 | $60 \pm 20$ | 0.83 |
| C6A/G13U | SpyRho555 | $40 \pm 4$ | 0.57 |

<sup>a</sup> $K_{\text{rel}} = K_{D, \text{WT+other dye}}/K_{D, \text{WT+SpyRho555}}$ or $K_{D, \text{mutant}}/K_{D, \text{WT+SpyRho555}}$
<sup>b</sup>N.D. not detected, N/A not applicable

For crystallization trials of RNA:fluorophore complexes, we chose to use 5(6)-TAMRA, a mixture of the 5- and 6-carboxy isomers. This dye is highly analogous to TMR, which RhoBAST has been shown to bind with high affinity (15). The only structural difference is the addition of a carboxylate group at the phenyl ring at the 5- or 6-position (Figure 1D). We quantified RhoBAST binding to 5- and 6-TAMRA and observed comparable affinities to SpyRho555 binding (Table 1, Supplemental Figure S1B). Unlike SpyRho555, which fluoresces only in the presence of the RhoBAST aptamer, the fluorescence of 5- and 6-TAMRA changes much less upon addition of RNA. As discussed below, there is a large shift in the absorption and emission wavelength of these dyes upon RNA binding. On a plate reader, with fixed excitation parameters optimized for the unbound dye, the wavelength shift gives rise to an apparent decrease in fluorescence upon RNA binding, which was used to measure K_D_ values. To validate our fluorescence binding data, we performed isothermal titration calorimetry (ITC) using WT RhoBAST and 5-TAMRA. Supporting our experimental setup, we obtained similar K_D_s from fluorescence binding (82 ± 4 nM, Table 1, Supplemental Figure S1B) and ITC (70 ± 20 nM, Supplemental Figure S2, Supplemental Table S3).

### RhoBAST structure contains a 4HJ with a network of interactions between J2/3 and L3

Crystals of the RhoBAST:5-TAMRA complex contained four protomers per asymmetric unit, and each protomer was built and refined individually. Despite both the 5- and 6-isomers of TAMRA being present in the crystallization conditions, electron density was only observed consistent with the 5-isomer of TAMRA bound to RNA (Supplemental Figure S3). Superimposition of the four protomers revealed good agreement with only small differences among structures both globally and locally (Supplemental Figure S4). Inspection of the lattice contacts of each protomer revealed significant intermolecular RNA:RNA interactions near the binding site in three of the molecules that may influence the details of the dyes’ interaction with the isolated aptamer (Supplemental Figure S5). Thus, we selected chain A, which does not contain lattice contacts near the binding site, for our analysis of the overall structure and binding site. Recently, another structure of RhoBAST in complex TMR-DN was independently determined (44), and the two structures are nearly identical at the global level (RMSD of 1.07 Å, Supplemental Figure S6).

Rather than the predicted three-way junction, the RhoBAST aptamer contains a 4HJ (Figure 2A-B). P1 and P3 coaxially stack on one another as do P2 and J2/3; the two coaxial stacks are oriented in an antiparallel configuration. This is the same configuration of helices found in the well-characterized hairpin ribozyme (45) and is a common arrangement among four-way junctions in RNA structures (46). This indicates that long-range tertiary interactions outside the junction in RhoBAST do not enforce an atypical arrangement of helices. At the junction, four Watson-Crick-Franklin (WCF) base pairs define this arrangement (Figure 2C). The U6-G13 base pair of P2 stacks with U14-A20 of J2/3 while G21-C42 of J2/3 stacks on U5-A43 of P1. The U5 residue, inserted during the engineering of RhoBAST to improve function (13), helps form a perfect junction whereas A43 was likely unpaired in the original SRB-2 aptamer.

**Figure 2.**
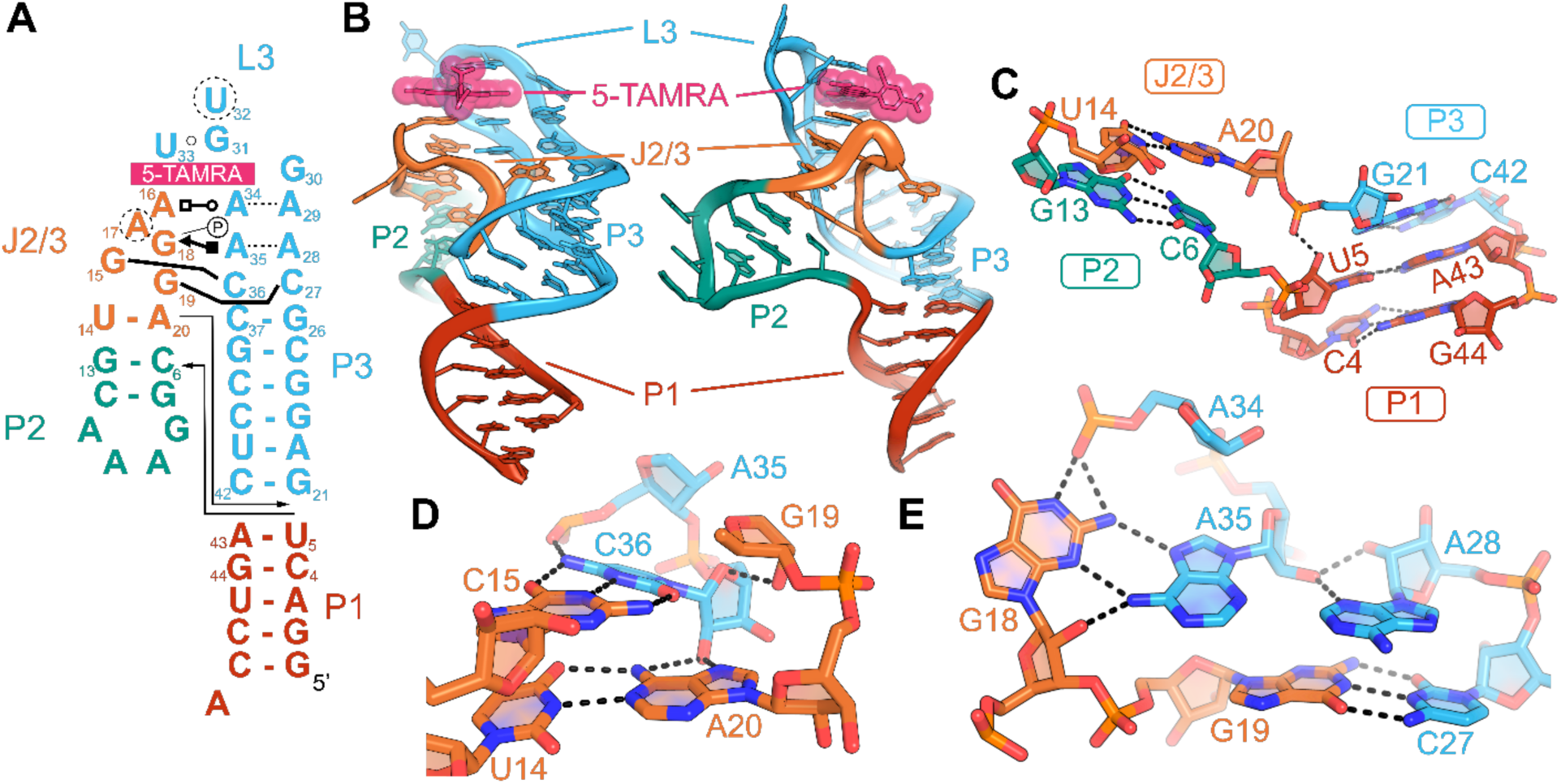
Secondary and tertiary structures of RhoBAST in complex with 5-TAMRA. (**A**) Secondary structure schematic of RhoBAST based on the crystal structure. (**B**) Global structure of RhoBAST shown as a cartoon from two angles. The ligand, 5-TAMRA, is shown in spheres. (**C**) Close-up view of nucleotides and interactions observed in the 4HJ shown as sticks. G21-C42 of P3 π-stack with U5-A43 of P1 while U14-A20 of J2/3 π-stacks with C6-G13 of P2. A single interaction between the 2′-hydroxyl of U5 (P1) and non-bridging phosphate oxygen of G21 (P3) can be seen in the junction. (**D**) One set of tertiary interactions between J2/3 and L3 shown as sticks. The U14-A20 (J2/3) WCF base pair flanking the junction makes an additional interaction with the 2′-hydroxyl of C36 (L3) through the Hoogsteen edge of A20 (J2/3). C36 (L3) forms a WCF base pair with G15 (J2/3), hydrogen bonds with the non-bridging phosphate oxygen of A35 (L3) with its N4 group, and interacts with the 2′-hydroxyl of G19 (J2/3) with its ribose oxygen. (**E**) Another set of tertiary interactions between J2/3 and L3 shown as sticks. G19 (J2/3) and C27 (L3) form a second WCF base pair between J2/3 and L3. A type II A-minor triple, G18-A35-A28, features a sheared G-A base pair with the sugar edge of G18 (J2/3) and Hoogsteen edge of A35 (L3) and A28 (L3) interacting through its N5 and 2′-hydroxyl with the 2′-hydroxyl of A35 (L3) to complete the triple. G18 (J2/3) makes an additional interaction with the non-bridging phosphate oxygen of A34 (L3).

To evaluate the tolerance of this junction to mutations, an important consideration for further engineering of RhoBAST, we mutated the G-C base pairs in P1 and P2 to a U•G mismatch or A-U base pair. Neither mutation (C4U, C6A/G13U) was deleterious to SpyRho555 binding (60 ± 20 nM and 40 ± 4 nM, respectively) and resulted in slight increases in affinity compared to WT (Table 1, Supplemental Figure S1C). Similarly, changing the A-U base pair of J2/3 to a G-C base pair results in no change in affinity from WT when the purine-pyrimidine identity is retained (U14C/A20G, 60 ± 20 nM, Table 1, Supplemental Figure S1D). However, flipping the purine-pyrimidine identity, U14G/A20C, results in a 2.6-fold weaker affinity than WT (180 ± 30 nM, Table 1, Supplemental Figure S1D), suggesting that π-stacking interactions have been optimized during selection of the original SRB-2 aptamer and doped re-selection of RhoBAST. Further supporting that this purine-pyrimidine positioning is important to RNA function, neither a purine at position 14 or a pyrimidine at position 20 was identified in the original selection of SRB-2 (26).

The only other observable interaction in the junction is a hydrogen bond between the 2′-hydroxyl of U5 in helix P1 and the non-bridging phosphate oxygen of G21 in helix P3 (Figure 2C). The observed electron density around this region does not support placement of any metal ions in the junction in any of the four protomers of our RhoBAST:5-TAMRA model. This contrasts the recently determined RhoBAST:TMR-DN structure (44) that contains two magnesium ions in the junction which the authors conclude are critical for holding J2/3 and P3 together. Our structure suggests otherwise, as we obtained an overall identical structure that does not support cation placement in the locations identified in the RhoBAST:TMR-DN structure. Consistent with our observation is the lack of metal cations in the crystal structure of the 4HJ-containing hairpin ribozyme (45). Further, prior work has demonstrated that 4HJs can fold in the absence of Mg^2+^ by using high concentrations of monovalent sodium ions (47). Evaluating the binding affinity of RhoBAST to SpyRho555 in high concentrations of monovalent cations without Mg^2+^ (1 M NaCl, 1 mM EDTA) reveals that the RNA is still able to fold and bind the dye with a K_D_ of 110 ± 20 nM (Supplemental Figure S7). These data suggest that while RhoBAST folds more efficiently in low concentrations of Mg^2+^, there are not specific sites in the aptamer occupied by Mg^2+^. Instead, a high ionic strength can facilitate folding and function of RhoBAST in the absence of magnesium ions.

Reinforcing the global architecture of the aptamer are a series of tertiary interactions between J2/3 and L3. Above the U14-A20 base pair proximal to the junction in J2/3 is a WCF pair between G15 of J2/3 and C36 of L3 (Figure 2D). This stabilizes J2/3 by extending the A-form like helix by a second base pair. This small helical element is further reinforced by interactions between the 2′-hydroxyl of C36 and the N6 and N7 of A20, the ribose oxygen of C36 and the 2′-hydroxyl of G19 in J2/3, and N4 of C36 with the non-bridging phosphate oxygen of A35.

A second WCF base pair connecting J2/3 and L3 is between G19 and C27 (Figure 2E). This base pair serves to cap the P3 helix and stacks with the A28-G30 purine stack that helps form the 5-TAMRA binding pocket. Reinforcing the importance of this pair’s role in supporting the ligand binding pocket, disruption of this base pair by mutating G19 to a cytidine completely ablates binding (Table 1, Supplemental Figure S1E). The compensatory mutation, G19C/C27G, rescues binding, albeit with an affinity 33-fold weaker than WT (Table 1, Supplemental Figure S1E). While this suggests that the observed WCF interaction between G19 and C27 in the crystal structure is occurring, swapping the purine-pyrimidine identities results in a suboptimal stacking interaction with the purine stack.

Further distal from the junction, a planar surface is formed by pairing between G18 of J2/3 and A35 of L3 that is further augmented by a type II A-minor triple with A28 of the purine stack (Figure 2E). The central pairing interaction is mediated by the sugar edge of G18 contacting the Hoogsteen face of A35. G18 makes an additional interaction through its WCF edge with the non-bridging phosphate oxygen of A34 in L3. The N3 and 2′-hydroxyl of A28 contact the 2′-hydroxyl of A35 in a type II A-minor motif. Together, this extensive network of interactions between J2/3 and L3 establishes the ligand binding pocket between the tips of these two loops.

### RhoBAST binding site stacks with ligand and interacts with its 3-position group

The terminal loops of J2/3 and L3 form the RhoBAST binding pocket to facilitate interactions with fluorophore ligands (Figure 3A). Stacking on the G18-A35-A28 triple is a set of three purine bases that form the floor of the ligand binding pocket. The core of this floor is formed via base pairing between the WCF face of A16 in J2/3 and the Hoogsteen face of A34 in L3 (Figure 3B). This pair is supported by interactions between the N6 of A16 and the 2′-hydroxyl group of U33 and by A29 of the purine stack forming a type-II A-minor triple with A34. The xanthene moiety of 5-TAMRA stacks on top of the A16-A34 pair, complemented by π-cation interactions between the partially positive *N, N*-dimethylamine groups at the 3′ and 6′ positions of 5-TAMRA and the adenine bases.

**Figure 3.**
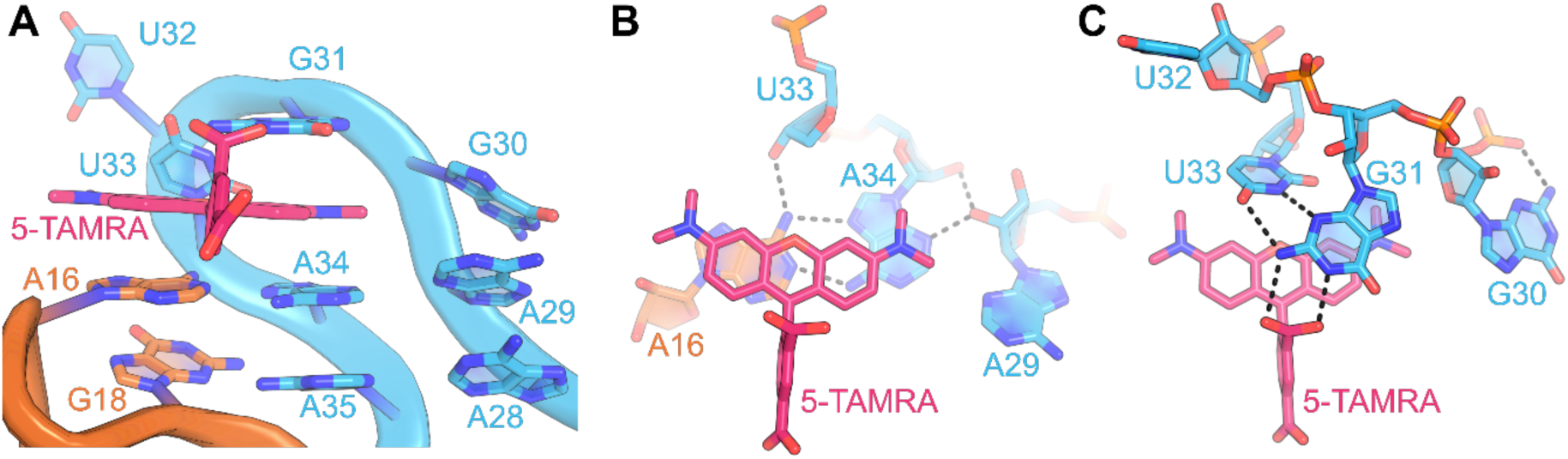
RhoBAST binding pocket with 5-TAMRA. (**A**) Overall view of RhoBAST binding pocket with the RNA shown as a cartoon and 5-TAMRA shown in sticks. 5-TAMRA stacks with a tri-adenine floor formed by nucleotides in J2/3 and L3. This tri-nucleotide interactions π-stacks directly on the base triple of G18-A35-A28. A purine stack is formed off to the side of the ligand by residues A28 through G30. G31 completes the π-stacking interactions with the ligand. (**B**) Tri-adenine floor of binding pocket represented as sticks. The adenine floor of the binding site contains a type II A-minor motif with the WCF edge of A16 (J2/3) interacting with the Hoogsteen edge of A34 (L3) and the 2′-hydroxyl of A29 (L3) contacting the sugar edge and 2′-hydroxyl of A34 (L3). N6 of A16 (J2/3) makes an additional interaction with the 2′-hydroxyl of U33 (L3). The xanthene core of 5-TAMRA π-stacks directly with A16 (J2/3) and A34 (L3) of the adenine floor. (**C**) Binding site interactions with 5-TAMRA shown as sticks. The WCF face of U33 contacts the sugar edge of G31 which π-stacks with the xanthene core and interacts directly with the ligand through its WCF face with the 3-carboxylate of 5-TAMRA. Adjacent to this interaction, G30 adopts the *syn* configuration and forms a hydrogen bond with its non-bridging phosphate oxygen.

On the other side of the bound 5-TAMRA, four nucleotides in L3 (G30-U33) form a turn structure that further encapsulates the dye. At the center of this turn, 5-TAMRA hydrogen bonds through its 3-carboxylate with N1 and N3 of G31 (Figure 3C). This guanine π-stacks above the xanthene with its sugar edge forming two hydrogen bonds with the WCF face of U33. To the side, G30 stacks with A29 as part of the purine stack, adopting a *syn* configuration that allows its N3 to hydrogen bond with one of its own non-bridging phosphate oxygens. In contrast to the RhoBAST:TMR-DN structure (44), the electron density around this nucleotide clearly supports the unambiguous assignment of the *syn* configuration. While the conformation of G30 in two protomers (B and D) may be influenced by lattice contacts, protomers A and C do not have neighboring lattice interactions which suggests that this is the solution conformation of the nucleotide (Supplemental Figure S8). Examination of the electron density and model of the RhoBAST:TMR-DN structure (PDB ID: 8JY0) (44) at the equivalent nucleotide position (G46) revealed poor local agreement (Supplemental Figure S9A-B). Adjusting G46 of chain B to the *syn* configuration produces a significantly better fit to the electron density without the need for two additional solvent molecules to fill density unoccupied by the guanine base (Supplemental Figure S9C-D). The *syn* configuration of G30 in our structure precludes any interaction with G31, a further difference in the details of the RNA structure surrounding the bound dye between the two models (44) (Supplemental Figure S9E-F).

The interpretation of the binding pocket electron density results in differing interactions between the RNA and the TMR moiety shared by ligands used in the RhoBAST:TMR-DN (44) and RhoBAST:5-TAMRA structures. The binding site of protomer B, the only protomer considered in the analysis of the TMR-DN structure, shows the equivalent G31 residue (G47) interacting with one side of the 3-carboxylate of TMR and the quencher moiety, DN (44) (Supplemental Figure S10A). However, the quencher group of the TMR-DN ligand in protomer B π-stacks with the TMR phenyl ring present in protomer F, dictating its position in the structure (Supplemental Figure S10B-C). Given both the higher resolution of the RhoBAST:5-TAMRA structure (2.10 Å versus 2.75 Å) and that protomer A in the 5-TAMRA structure does not have lattice contacts adjacent to the dye binding pocket, we believe that our model more accurately reflects the solution conformation of RhoBAST bound to dye.

### RhoBAST in complex with MaP555 is globally identical to RhoBAST:5-TAMRA structure

SpyRho555 contains an *N, N*-dimethylsulfamide group at its 3-position instead of the carboxylate present in 5-TAMRA. To determine how RhoBAST interacts with the 3-position moiety involved in the spirocyclization of SpyRho555, we co-crystallized RhoBAST with both SpyRho555 and the analogous compound, MaP555. MaP555 shares the same core as SpyRho555 with the only difference being the addition of a 6-carboxylate group that is present in 6-TAMRA (Figure 1D, F). The affinity of WT RhoBAST for MaP555 is 120 ± 20 nM (Table 1, Supplemental Figure S1F) compared to SpyRho555 binding (70 ± 20 nM, Table 1), demonstrating that RhoBAST accommodates both ligands in a similar manner. While we obtained crystals of RhoBAST in the presence of both SpyRho555 and MaP555, only the crystals containing MaP555 diffracted sufficiently for structural determination.

The crystal of the RhoBAST:MaP555 complex contained two protomers in the asymmetric unit and neither exhibited significant lattice contacts near the binding site (Supplemental Figure S11). However, the ligand density in protomer B was too sparse to confidently place MaP555, so we chose not to place a ligand in this molecule in the final structural model. We caution against interpretations that protomer B represents the apo conformation as the data quality in this region is poor and does not support this conclusion. In contrast, protomer A has well-defined density for the ligand (Supplemental Figure S12) and displays better resolved RNA features around the binding pocket. Thus, this protomer is considered representative of the bound RNA:dye structure.

Globally, this RhoBAST:MaP555 structure is identical to the RhoBAST:5-TAMRA structure with an all atom RMSD of 1.9 Å (Supplemental Figure S13). The one significant difference between the RNA structures is observed in the GAAA tetraloop of P2. In the RhoBAST:MaP555 structure, this loop adopts the canonical configuration where the first and fourth nucleotides form a sheared G-A base pair and the last three adenosine residues stack on top of one another (48). The RhoBAST:5-TAMRA structure displays an unusual conformation where the fourth residue is displaced outward and no longer π-stacks with the third residue of the loop or forms a sheared pair with the first position guanine. Examining the lattice contacts occurring in this region, the P2 tetraloop of protomer A extensively interacts with the P2 tetraloop of protomer D (Supplemental Figure S14). Three base triples occur between the GAAA tetraloop of protomer A and protomer D. The first residue of the protomer A tetraloop, G8, interacts through its sugar edge with the WCF face of the third residue, A10, in the protomer D tetraloop which further contacts N6 of A11 in protomer A through its sugar edge (Supplemental Figure S14B). The second and third residues of the protomer A tetraloop engage in base triples with residues in protomer D (Supplemental Figure S14C-D). This previously unobserved “head-to-head” interaction between two GAAA tetraloops reveals yet another way that this loop motif can facilitate long-range RNA:RNA contacts.

### RhoBAST interacts with 3-position group of MaP555

The MaP555 dye is located at the same binding site as 5-TAMRA with nearly identical positioning (Figure 4A, Supplemental Figure S15A). The xanthene groups of both ligands stack over A16-A34 of the adenine floor and superimpose well with each other although the core of MaP555 is rotated slightly out of the binding site compared to 5-TAMRA (buried surface area: 327 Å^2^ for MaP555 and 336 Å^2^ for 5-TAMRA, Figure 4B, Supplemental Figure S15B). The phenyl group of MaP555 is rotated at a steeper angle relative to the core of the dye than 5-TAMRA (11° difference, Supplemental Figure S15C-D), orienting the 3-position *N, N*-dimethylsulfamide for interactions with G31. Like the 5-TAMRA structure, G31 hydrogen bonds through its WCF face with the 3-position of MaP555 (Figure 4C). N1 and N2 of G31 engage with the amide carbonyl and sulfur oxygen of MaP555. The N3 of U33 still engages with N3 of G31, but the other hydrogen bond observed between the 4-carbonyl of U33 and N2 of G31 is lost (3.1 Å distance in 5-TAMRA structure vs 3.9 Å distance in MaP555 structure). The 3-position moiety of MaP555 is conserved in SpyRho555, so this structure should represent how RhoBAST interacts with the SpyRho555. The 6-carboxylate of the phenyl ring, the only structural difference between MaP555 and SpyRho555, does not interact with the RNA and faces solvent.

**Figure 4.**
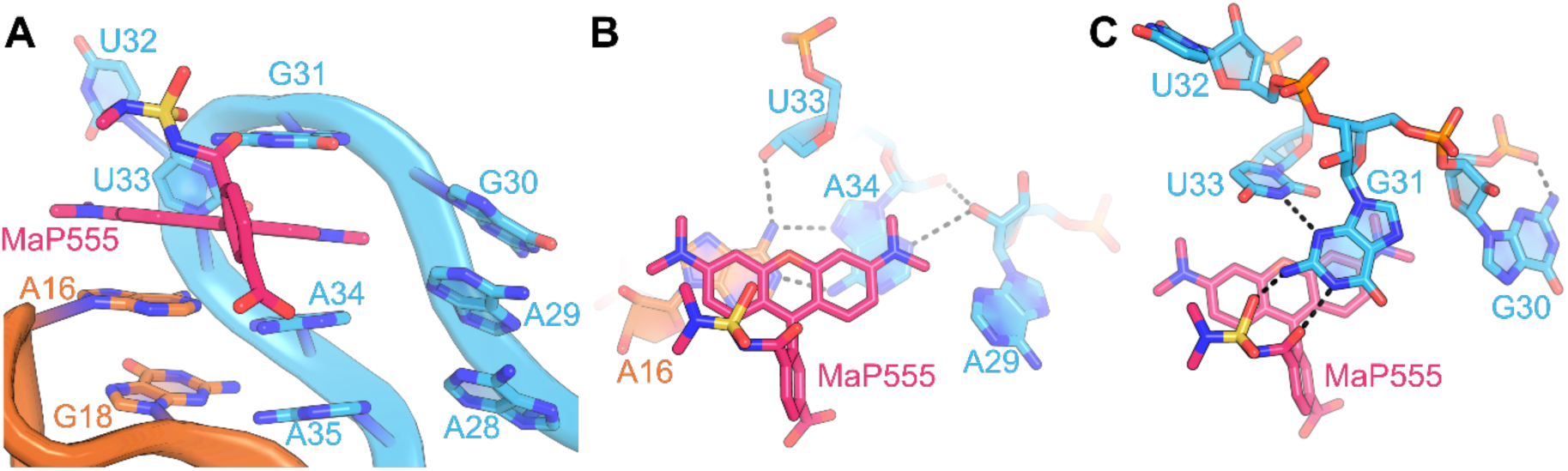
RhoBAST binding pocket with the SpyRho555 analogue, MaP555. (**A**) Overall view of the RhoBAST binding pocket as a cartoon with MaP555 shown as sticks. MaP555 binds in the same location as 5-TAMRA, sandwiched between the tri-adenine floor composed of nucleotides from J2/3 and L3 and G31 of L3. (**B**) Adenine floor of the binding site shown as sticks. The tri-adenine floor, A16-A34-A29, is a type II A-minor triple that forms the basis of the ligand binding site. The xanthene core of MaP555 π-stacks with A16 (J2/3) and A34 (L3). The interactions observed in the RhoBAST:MaP555 structure are identical to the RhoBAST:5-TAMRA structure. (**C**) Interactions between the RNA and ligand are shown as sticks. The WCF of U33 interacts with the sugar edge of G31 identical to the interactions observed in the RhoBAST:5-TAMRA structure. However, the interaction between the 4-carbonyl of U33 and N2 of G31 is lost. The WCF face of G31 interacts with the 3-position *N, N*-dimethylsulfamide moiety of MaP555, specifically interacting with the sulfur oxygen and the amide carbonyl groups.

This structure implicates G31 as important for SpyRho555 binding through its interactions with the 3-position group. Mutating G31 to an adenine, G31A, drastically reduces the ability of RhoBAST to stabilize the open form of SpyRho555 and weakens the affinity through fluorescence binding measurements by greater than 430-fold (Table 1, Supplemental Figures S1G). A G31C mutant further weakens the affinity to greater than 640-fold (Table 1, Supplemental Figure S1G). Because these binding measurements rely on the ability of RhoBAST to turn-on SpyRho555, we cannot determine if the observed weaker affinity is solely due to a lack of binding or a loss in the RNA’s ability to stabilize the fluorescent configuration of SpyRho555 through interactions between G31 and the 3-position moiety of the dye. Regardless, the interactions between G31 and the 3-position of its ligands as seen in the 5-TAMRA and MaP555 structures are crucial for conferring the fluorogenic capabilities of RhoBAST and provide an explanation as to how RhoBAST stabilizes the open configuration of SpyRho555 to increase fluorescence using the ring-opening mechanism.

### Chemical footprinting of RhoBAST indicates a preorganized aptamer

To examine the effect of dye binding on RNA structure in solution, we used SHAPE chemical probing. This approach relies on the ability of an acid anhydride (*N*-methylisatoic anhydride, NMIA) to react with 2′-hydroxyl groups in conformationally flexible regions of the RNA that can be read out as a reverse transcriptase stop in a sequencing reaction (39). The reactivity pattern observed in the absence of ligand agrees well with the RhoBAST crystal structure (Figure 5A, Supplemental Figure S16). The terminal loops, L2 and L3, display greater reactivity than the surrounding nucleotides that are engaged in helices, P2 and P3, respectively. For example, in J2/3 in the crystal structure, U14-A20 is observed making a WCF base pair, consistent with the low reactivity observed for these nucleotide positions. In contrast, A17 of J2/3 is flipped out into solution in the crystal structure and displays greater reactivity with NMIA. In L3, most nucleotides show low reactivity, suggesting a rigid structure; reactivity of the single flipped out nucleotide (U32) is obscured by an intrinsic reverse transcriptase stop.

**Figure 5.**
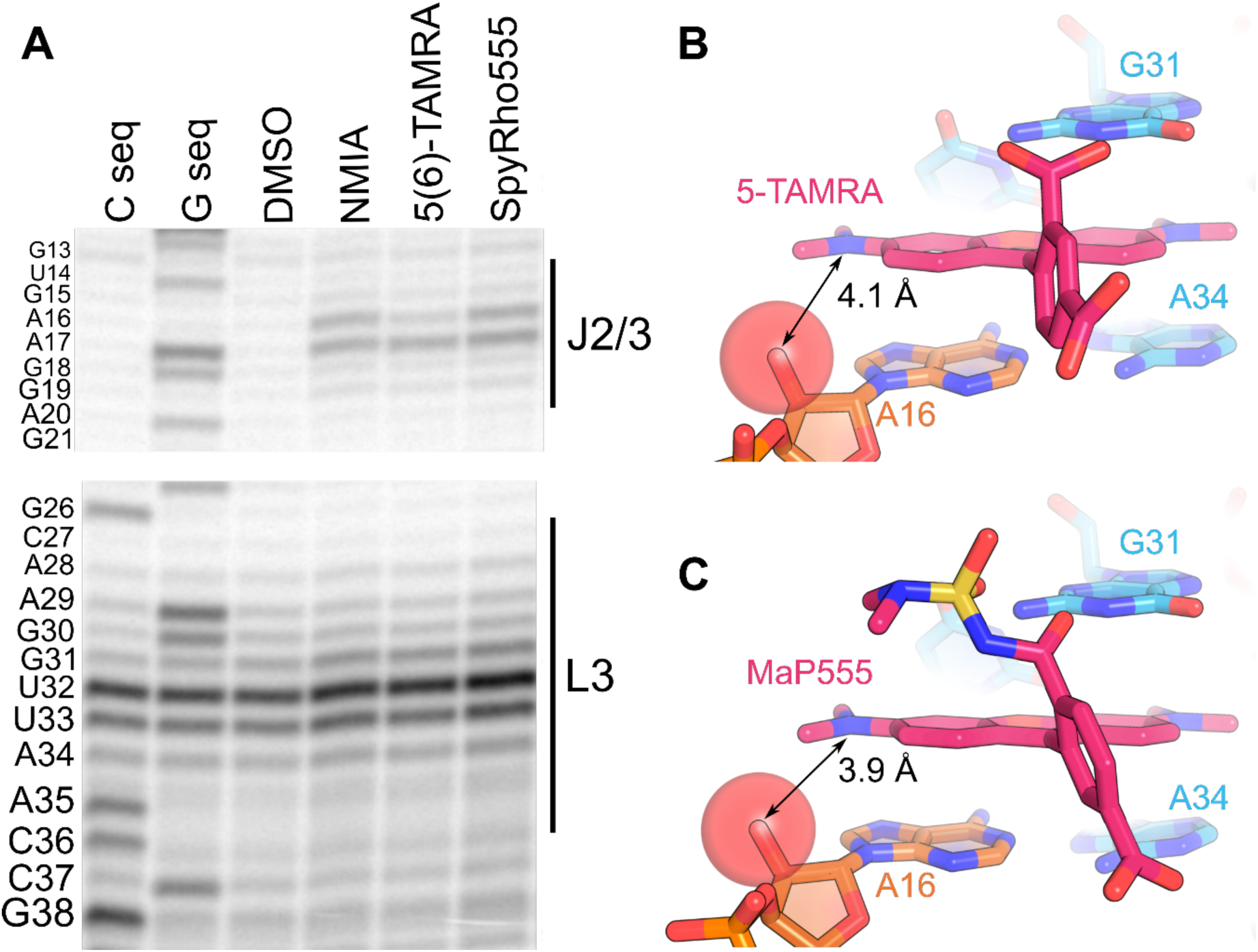
Chemical footprinting of the RhoBAST aptamer in the absence and presence of ligand. (**A**) SHAPE sequencing gel of RhoBAST in the absence of ligand or with 10 µM of 5(6)-TAMRA or SpyRho555. (B) Location of A16 and proximity to 5-TAMRA shown as sticks. A16 of J2/3 is part of the tri-adenine floor of the binding pocket and lies within proximity to the ligand. The 2′-hydroxyl of A16 (shown as a sphere) is 4.1 Å away from one of the amine groups of the xanthene ring. (**C**) Location of A16 and proximity to MaP555 shown as sticks. A16 (J2/3) adopts a similar position in the RhoBAST:MaP555 structure to the positioning seen in the RhoBAST:5-TAMRA structure. The distance between the amine substituent of the xanthene in MaP555 and 2′-hydroxyl of A16 (shown as a sphere) is 3.9 Å. The bulkier 3-position group of MaP555 is positioned closer to A16 than the 3-carboxylate of 5-TAMRA.

Consistent with previous chemical footprinting analysis of the RhoBAST aptamer (40), the NMIA reactivity pattern of the RNA does not significantly change in the presence of either 5(6)-TAMRA or SpyRho555. However, we observed a single chemical protection corresponding to the 2′-hydroxyl group of A16 only in the presence of 5(6)-TAMRA (Figure 5A). A16, which resides in J2/3, is one of the three nucleotides that forms the adenine floor of the ligand binding pocket with A34 and A29 of L3. The 2′-hydroxyl of A16 in the RhoBAST:5-TAMRA structure is directed towards the ligand and is not observed interacting with any other groups (Figure 5B). The position of this 2′-hydroxyl is the same in the RhoBAST:MaP555 structure, which should represent the RNA bound to SpyRho555, but no change in reactivity is observed in the presence of SpyRho555 (Figure 5C). The electrophilicity of a 2′-hydroxyl group and thus its reactivity towards NMIA is highly influenced by the local chemical environment (39). The most significant chemical difference between SpyRho555 and 5(6)-TAMRA is the identity of the 3-position moiety which may influence the local electrostatic environment enough to generate different reactivity patterns between the two bound complexes at the 2′-hydroxyl of A16. Despite this single reactivity difference between the two dyes, the chemical probing data strongly supports that the RNA undergoes little, if any, conformational change upon binding rhodamine dyes. A pre-formed, rigid dye binding pocket also suggests that TMR derivatives carrying bulkier amine substituents on the xanthene ring or silicon rhodamine dyes would be strongly rejected by this RNA, but bulky modifications of the 3-position would be tolerated.

### Binding to RhoBAST alters the spectral properties of the rhodamine dyes

In aqueous solution, 5-TAMRA, 6-TAMRA, and MaP555 primarily exist in their open zwitterionic forms whereas SpyRho555 remains in the closed spirolactam form. RhoBAST stabilizes the open form of SpyRho555, such that upon binding a significant increase in absorbance and fluorescence emission is observed for the dye due a shift in the equilibrium from the closed colorless spirolactam to the open form (15).

In contrast, for the dyes with the equilibrium shifted toward the open zwitterionic form in aqueous solution (MaP555, 5-TAMRA, and 6-TAMRA), RNA binding induces a slight decrease in absorbance, an increase in fluorescence, and a bathochromic shift (Figure 6A-C). The extinction coefficients decrease between 18-27% (Figure 6D) while the fluorescence emission increases between 1.6- and 1.9-fold. Finally, all tested dyes exhibit a bathochromic shift in the absorption maximum of up to 12 nm and emission maximum of up to 10 nm upon RNA binding (Figure 6D). The changes in electronic structure are likely due to extended electron delocalization resulting from base π-stacking interactions between the dye and RhoBAST in addition to rigidification of the dye upon RNA binding that alters non-radiative decay pathways. Changes in the polarity and ionic conditions surrounding the dye could also affect both absorption and emission. Overall, the changes in electronic structure for MaP555, 5-TAMRA, and 6-TAMRA are similar to what has been reported for TMR (15).

**Figure 6.**
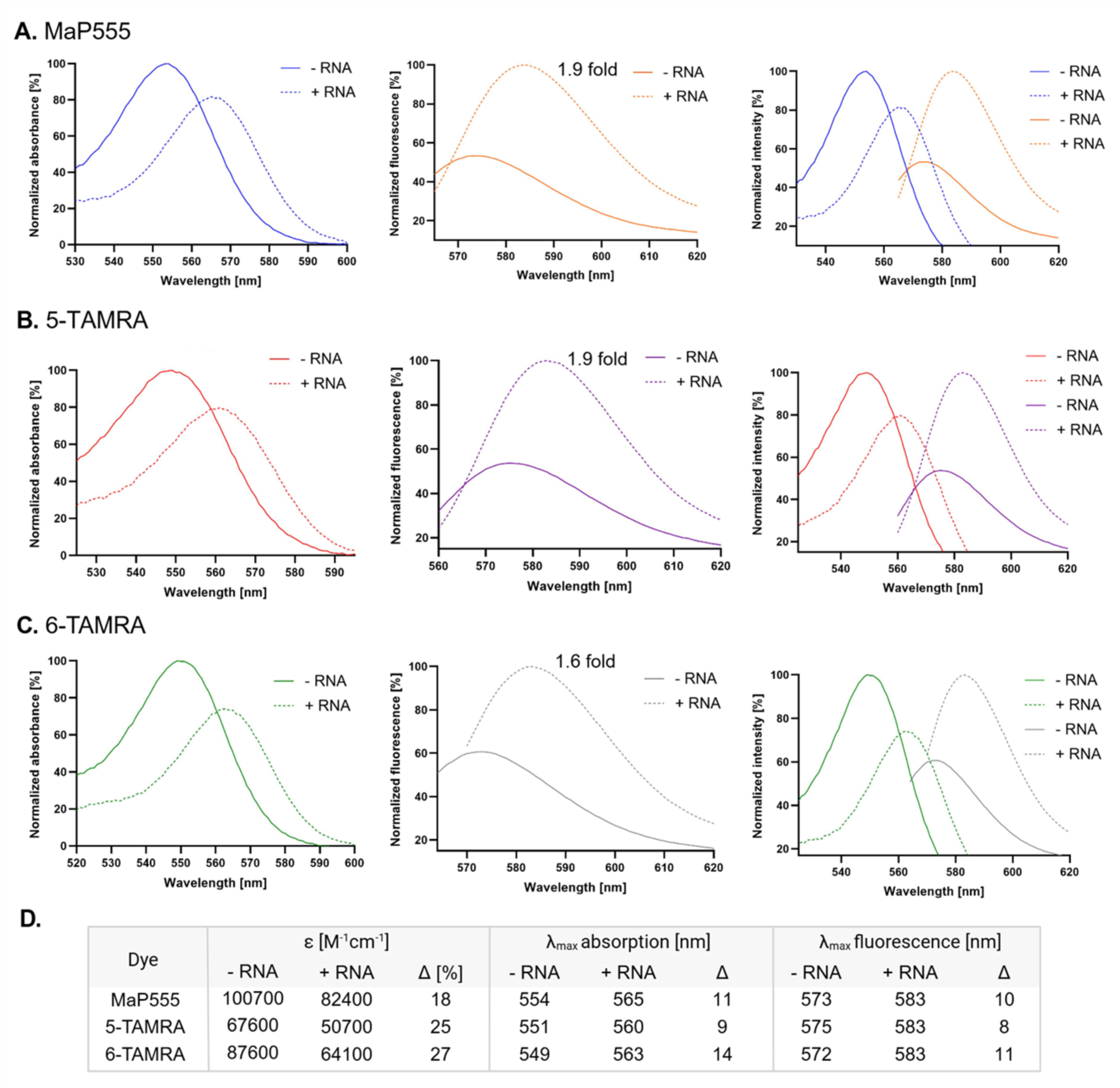
Spectral properties of rhodamine dyes in the presence and absence of RhoBAST. (**A**) From left to right: absorption spectrum, fluorescence emission spectrum, absorption and fluorescence emission spectra of MaP555 with and without RhoBAST. (**B**) From left to right: absorption spectrum, fluorescence emission spectrum, excitation and fluorescence emission spectra of 5-TAMRA with and without RhoBAST. (C) From left to right: absorption spectrum, fluorescence emission spectrum, excitation and fluorescence emission spectra of 6-TAMRA with and without RhoBAST. (**D**) Table summarizing the spectral changes observed when the dye binds to RhoBAST. Δ represents a change observed in the spectrum. Because of the bathochromic shift in absorbance, samples were excited at the isosbestic point to measure fluorescence changes upon RNA binding.

## DISCUSSION

Here, we have determined co-crystal structures of the RhoBAST aptamer bound to two rhodamine dyes: 5-TAMRA and an analogue of SpyRho555, MaP555. RhoBAST is organized by a 4HJ that enables J2/3 and L3 to form a network of tertiary interactions that establishes the peripheral dye-binding pocket. The xanthene core of 5-TAMRA is sandwiched between a purine floor comprised of nucleotides from J2/3 and L3 and G31 in L3. This interaction is mediated primarily through π-stacking. Hydrogen bonding interactions occur between the 3-position of the ligand with the WCF edge of G31. In the RhoBAST:MaP555 structure, the WCF face of G31 interacts with a non-bridging sulfur oxygen and amide carbonyl of the 3-position group that is structurally identical to the moiety present in SpyRho555. Mutation of G31 drastically weakens binding affinity implicating this RNA:ligand interaction as vital for RhoBAST to effectively interact with the spirocyclic dye SpyRho555.

RhoBAST represents one of the three current fluorogenic aptamers that shifts the equilibrium of dyes from the closed to open state to enable fluorescent turn-on. SiRA binds a silicon rhodamine dye structurally analogous to TMR that contains a silicon atom at the 10′-position of the xanthene ring (49) while the BeCA aptamer uses a ratiometric dye composed of benzopyrylium and coumarin groups (50). Because the sequences and predicted secondary structures of SiRA and BeCA differ from RhoBAST, the crystal structure of RhoBAST cannot be used to infer structural features of these RNAs. It remains unknown if SiRA and BeCA stabilize the open configuration of their cognate ligands through interactions with the 3-position group responsible for spirocyclization, like RhoBAST, or through other currently unknown interactions. It is also unclear if RhoBAST provides a unique local environment to facilitate ring-opening fluorescence or if any interactions with the 3-position of group of SpyRho555 are sufficient to induce fluorescent turn-on. The structure of another rhodamine-binding aptamer, TMR3, was solved in the presence of 5-TAMRA, revealing similar interactions to RhoBAST with the 3-carboxylate of 5-TAMRA interacting with two guanine bases and the dominant binding interactions occurring through π-stacking (51). In contrast to RhoBAST slightly increasing the fluorescence of 5-TAMRA upon binding, TMR3 significantly quenches the fluorescence of 5-TAMRA (51), indicating very different local electronic environments within their respective binding pockets.

The structural core of the RhoBAST aptamer is organized by a 4HJ bringing together distal RNA elements to create the ligand binding site, an architectural theme observed in other functional RNAs. For example, the hairpin ribozyme, which contains a 4HJ in the antiparallel helical arrangement like RhoBAST, hosts its binding pocket at the interface of two inter-helical loops (45). This 4HJ enables efficient folding of the hairpin ribozyme in micromolar concentrations of Mg^2+^ while folding in the absence of this core structural component increases the need for Mg^2+^ by three orders of magnitude (52). Another *in vitro* selected fluorogenic aptamer using a four-way junction for structural organization is DIR2, which employs restriction of TICT to increase dye fluorescence (53). Similar to the architecture of RhoBAST, DIR2 uses tertiary interactions between two terminal loops to form a binding pocket that sandwiches the planar fluorophore between a tri-nucleotide floor and a single purine (53). However, unlike RhoBAST and the hairpin ribozyme, the DIR2 junction contains a single unpaired nucleotide and adopts the parallel helical configuration (53). DIR2 displays weaker affinity to its target dyes (high nanomolar to low micromolar range) and a significantly higher Mg^2+^ requirement (6-8 mM) than RhoBAST (54). The poorer binding and properties of DIR2 likely stem from a critical difference in the organization of its four-way junction.

The strengths imparted to the RhoBAST aptamer by its robust core architecture have the potential to be leveraged to create new aptamers using “scaffolded” *in vitro* selection. Rather than deep selection in which the starting RNA pool is fully randomized, scaffolded selection only randomizes a small pocket in the context of a larger folded RNA architecture (55). Using a three-way junction derived from the purine riboswitch as the scaffold (56), this approach has yielded novel aptamers that bind a variety of small molecules, including fluorophores (55, 57, 58). This scaffold enables selection of a binding pocket embedded within the junction while distal loop-loop interactions are part of the fixed architectural scaffold. The RhoBAST aptamer provides an alternative scaffold that would flip the binding pocket from a junction core to a site peripheral to the junction, enabling the strongest driver of folding to be retained as part of the scaffold. In this selection, the initial library would be fully embedded in the distal loops of J2/3 and L3, leaving the 4HJ and associated tertiary interactions to promote RNA architecture. This approach may provide an avenue for the selection of aptamers that bind to rhodamine dyes with superior performance to SpyRho555 that are rejected by RhoBAST.

While the MaP555 structure reveals how the RNA stabilizes the open form of SpyRho555, it does not shed light on the mechanism by which the RNA binds free dye. If the RNA uses a conformational selection mechanism, the aptamer interacts solely with the open form of SpyRho555, and the observed increase in bulk fluorescence is achieved by driving the open-closed equilibrium of the free dye towards the open state (Figure 7A). In contrast, an induced fit mechanism would allow for initial binding to the closed form of SpyRho555, and the RNA promoting conversion to the open form to increase fluorescence (Figure 7A). A docking analysis suggests that RhoBAST can bind the closed form of SpyRho555 where the xanthene core engages with the purine floor and has similar binding surface areas to MaP555 in the crystal structure (Figure 7B-C, Supplemental Figures 17-18, Supplemental Table S4). The top ranked docking poses of closed SpyRho555 have different orientations for the 3-position group: pointed towards G31 as observed in the crystal structures or facing away from G31 and interacting with the purine floor (Figure 7B-C). A proposed metal-mediated ring-opening mechanism for spirolactam rhodamines positions a cation proximal to the 3-moiety of the dye and further stabilizes the open form (21, 22) (Figure 7D). The configuration of G31 observed in the RhoBAST:MaP555 structure is not dissimilar to the proposed interactions that allow for metal-mediated ring-opening and stabilization. If RhoBAST binds the closed form of SpyRho555, G31 might facilitate ring-opening through interactions with the *N, N*-dimethylsulfamide group. While our analysis cannot rule out either binding model, our structure raises the intriguing possibility that the RNA may play dual roles in promoting ring-opening and stabilizing the open state.

**Figure 7.**
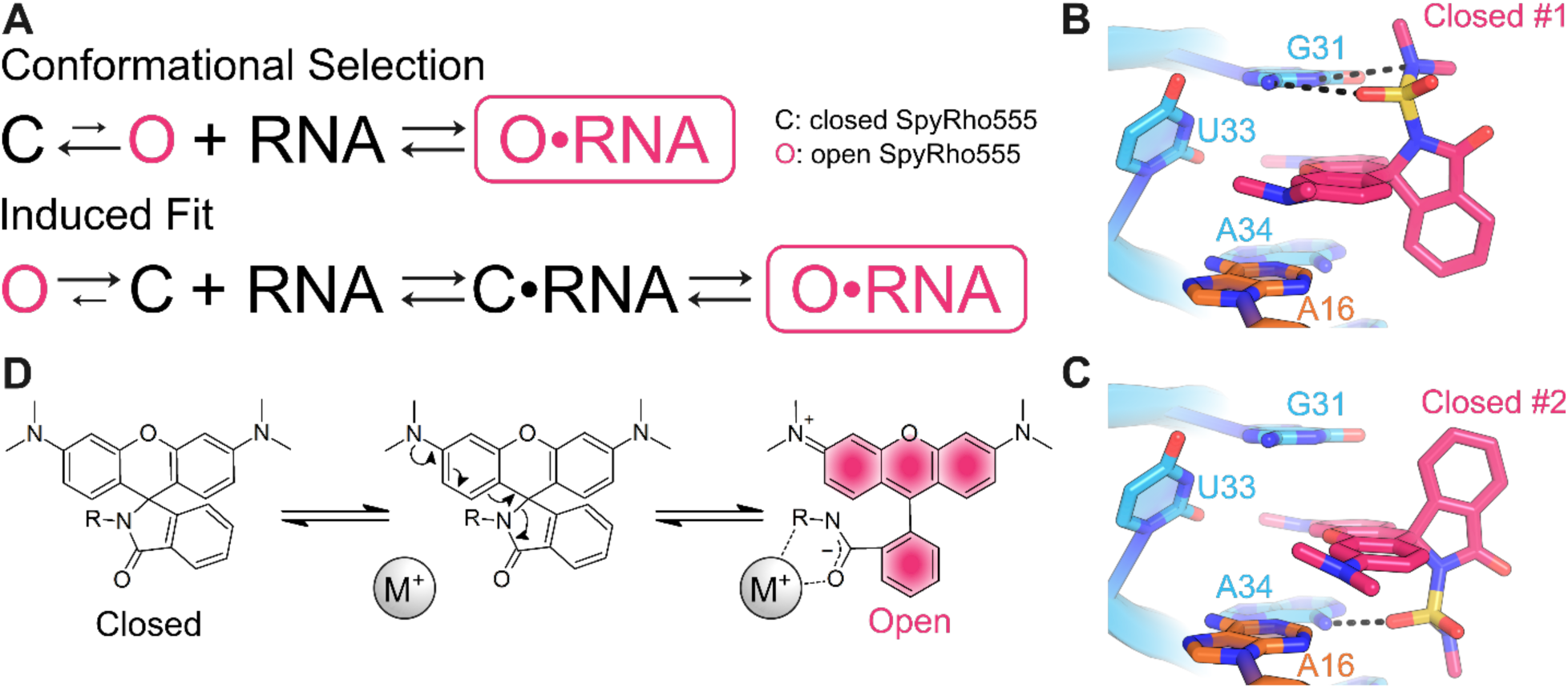
Two models for RhoBAST engaging with SpyRho555. (**A**) Schematic of conformational selection and induced fit models. (**B**) Top-ranked docking pose of closed SpyRho555. The 3-position group is facing towards G31 as observed in the crystal structures. (**C**) The second-ranked top docking pose of closed SpyRho555. The 3-position group is facing away from G31 and instead is predicted to interact with A34 of the tri-adenine floor of the binding pocket. (**D**) Proposed metal-mediated ring-opening mechanism.

## Data availability

Atomic coordinates and structure factor amplitudes have been deposited into the Protein Data Bank (PDB) database under accession codes 9BUN (RhoBAST:5-TAMRA) and 9DXL (RhoBAST:MaP555).

## Supplementary data

Supplementary Data are available at NAR Online.

## Acknowledgements

We thank Dr. Jay Nix and the staff of beamline 8.2.2. of the Advanced Light Source, Lawrence Berkeley National Laboratory for their support with remote crystallographic data collection. Beamline 8.2.2. of the Advanced Light Source, a DOE Office of Science User Facility under Contract No. DE-AC02-05CH11231, is supported in part by the ALS-ENABLE program funded by the National Institutes of Health, National Institute of General Medical Sciences, grant P30 GM124169-01. We thank the Macromolecular X-ray Crystallography Core (RRID:SCR_019310) at the University of Colorado Boulder for crystallographic data collection and the Shared Instruments Pool (RRID: SCR_018986) of the Department of Biochemistry at the University of Colorado Boulder for the use of the QM-6 PTI fluorescence spectrometer and the Typhoon 5. The Typhoon 5 is funded by NIH Shared Instrumentation Grant S10OD034218-01. We thank Dr. Annette Erbse for her assistance with the resources utilized at the University of Colorado Boulder. We thank Logan McCoy for generating RNA for footprinting experiments and assisting with fluorimeter measurements.

## Author contributions

S.H.S. conducted RNA crystallization, collected diffraction data, and solved the crystal structures. S.H.S. performed binding measurements. A.J.W. obtained extinction coefficients and spectra. A.J.W. synthesized and purified SpyRho555. S.R.L. performed footprinting experiments. L.T.O. performed docking analysis. S.H.S., A.J.W., A.E.P., and R.T.B. wrote the manuscript. All authors contributed to reviewing the data and editing the manuscript.

