## Supplemental Information for "Structural basis for ring-opening fluorescence by the RhoBAST RNA aptamer"

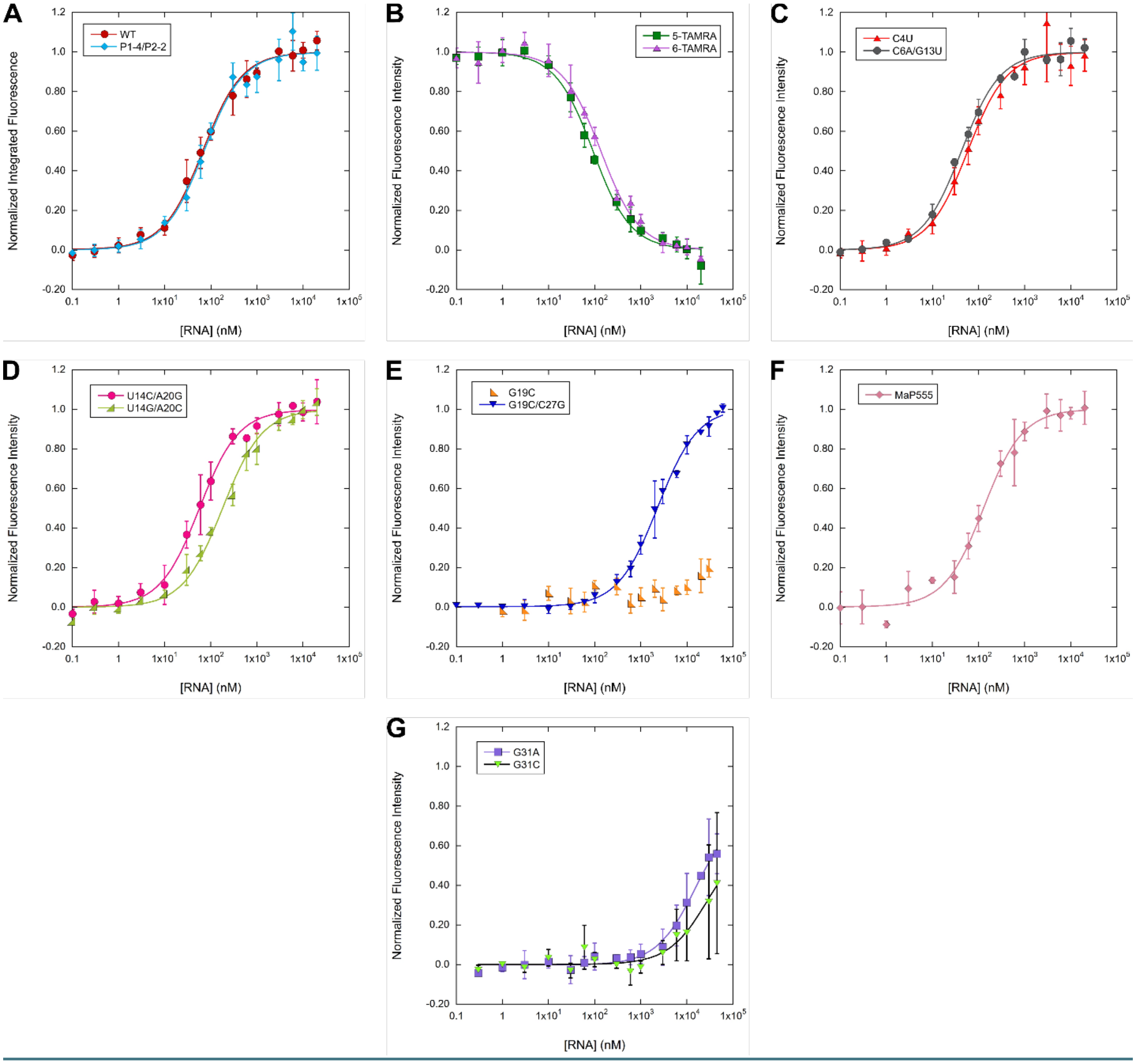

**Supplemental Figure S1.** Normalized fluorescence binding curves of wildtype (WT) RhoBAST, RhoBAST mutants, and dyes. (**A**) WT RhoBAST binding SpyRho555 (red) and P1-4/P2-2 binding SpyRho555 (blue). (**B**) WT RhoBAST binding 5-TAMRA (green) or 6-TAMRA (purple). (**C**) C4U (red) or C6A/G13U (gray) binding SpyRho555. (**D**) U14C/A20G (pink) or U14G/A20C (green) binding SpyRho555. (**E**) G19C (orange) or G19C/C27G (blue) binding SpyRho555. (**F**) WT RhoBAST binding MaP555 (pink). (**G**) G31A (purple) or G31C (green) binding SpyRho555. Data are shown as mean and standard deviation (s.d.) from independent measurements (n = 3-4).

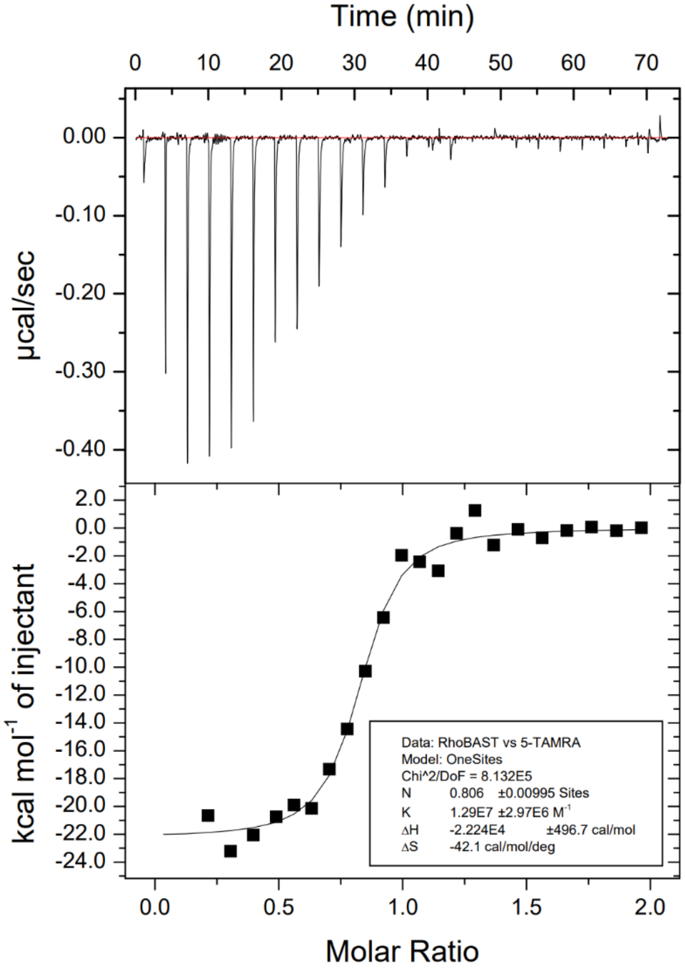

**Supplemental Figure S2.** Isothermal titration calorimetry binding isotherm of WT RhoBAST binding to 5-TAMRA. RNA was prepared as described in Methods and dialyzed using 6-8 kDa MWCO dialysis tubing (Spectra/Por) against 500 mL of 1x Aptamer Selection Buffer (20 mM HEPES pH 7.4, 125 mM KCl, 1 mM $MgCl_2$) for 12 hours at 4 °C. 5-TAMRA solid was brought up in the same buffer used to dialyze the RNA. Measurements were taken at 25 °C with 5-TAMRA injected into RNA using a MicroCal iTC200 (Malvern). Data were fit in Origin using the single site model. 5-TAMRA injected into buffer was used for background subtraction, and measurements were performed in triplicate. A representative binding isotherm is shown above.

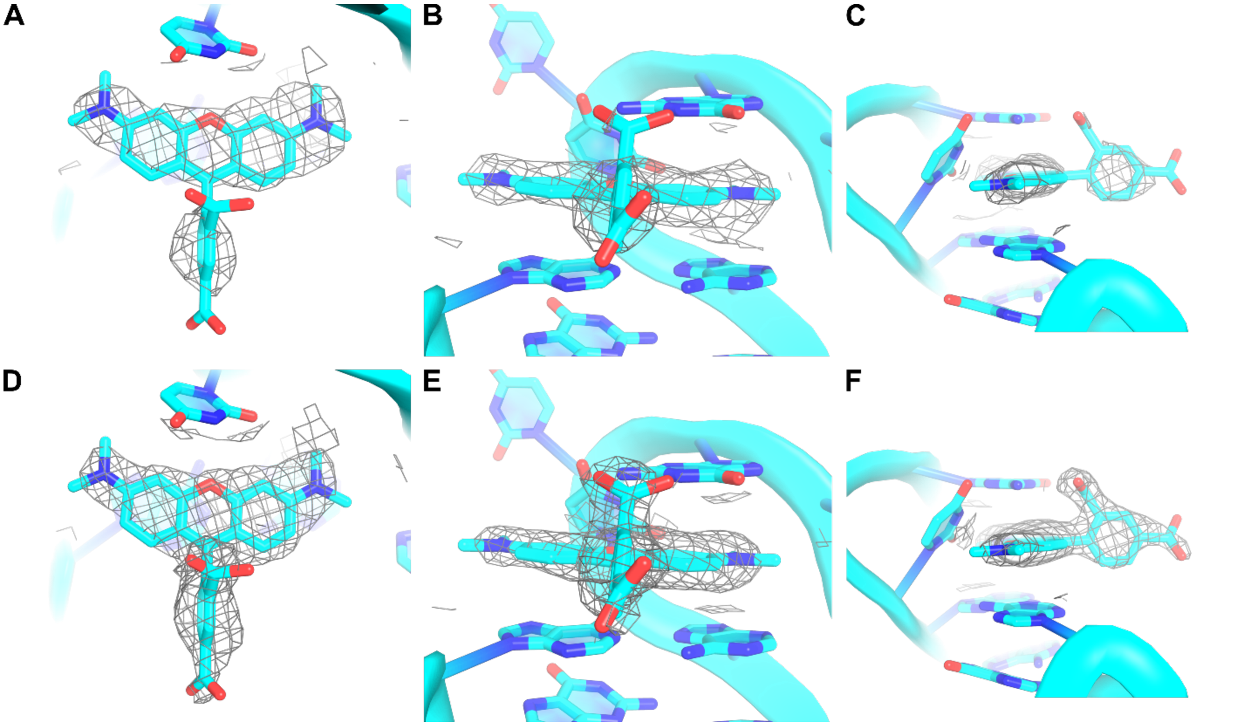

**Supplemental Figure S3.** Electron density supporting placement of 5-TAMRA in RhoBAST:5-TAMRA structure (PDB ID: 9BUN). All $2|F_o| - |F_c|$ maps are carved 2 Å around 5-TAMRA and shown at 1 σ. (**A**) Top-down view of 5-TAMRA with density modified map obtained directly from Phaser (1). The G31 base is removed for clarity. (**B**) View of 5-TAMRA in binding pocket with density modified map obtained directly from Phaser (1). (**C**) Sideview of 5-TAMRA phenyl ring with density modified map obtained directly from Phaser (1). (**D**) Top-down view of 5-TAMRA with simulated-annealing composite map (2). The G31 base is removed for clarity. (**E**) View of 5-TAMRA in binding pocket with simulated-annealing composite omit map (2). (**F**) Sideview of 5-TAMRA phenyl ring with simulated-annealing composite omit map (2).

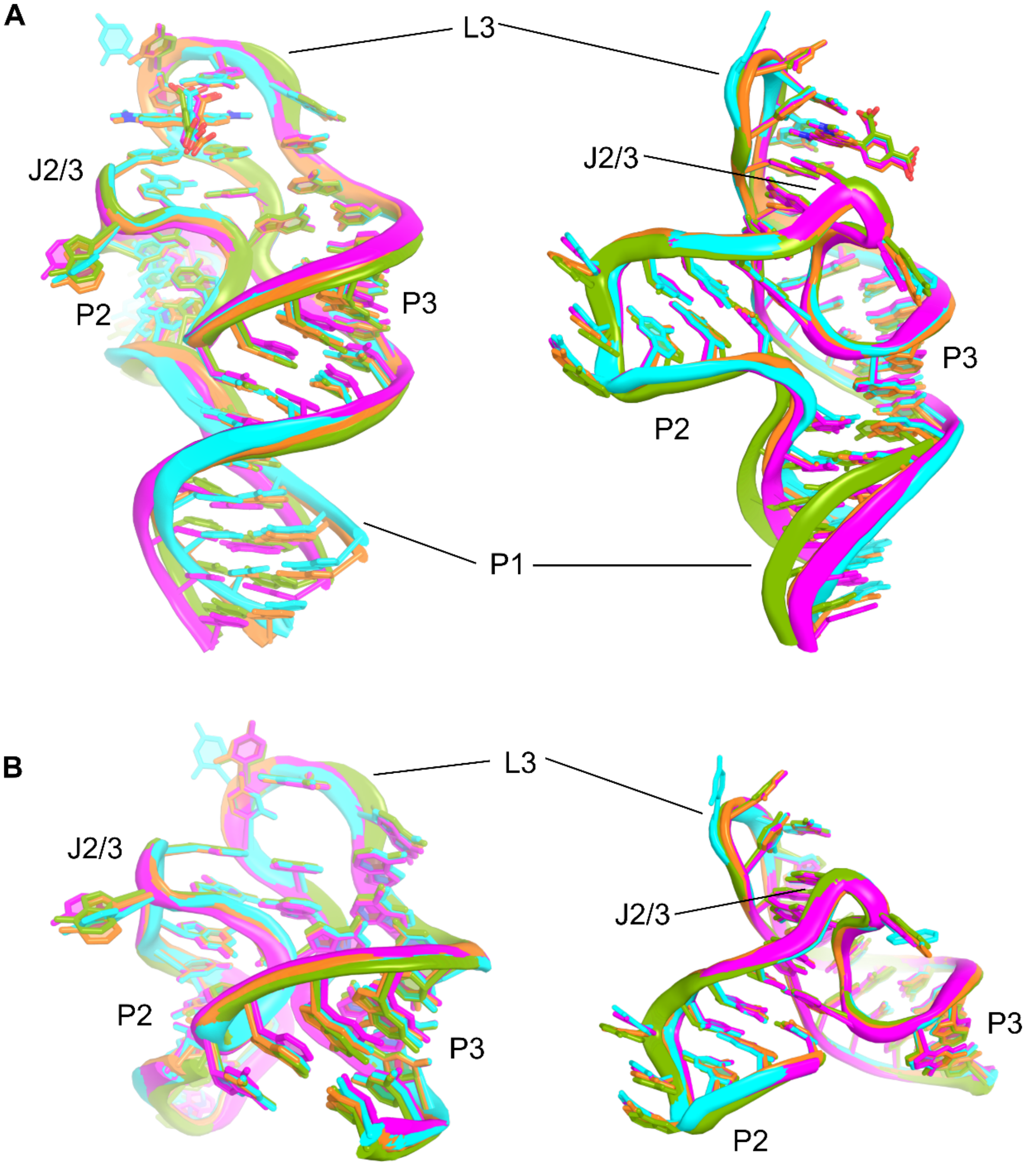

**Supplemental Figure S4.** Superimposition of all protomers in RhoBAST:5-TAMRA co-crystal structure. The color scheme is as follows: protomer A is cyan, protomer B is orange, protomer C is green, and protomer D is magenta. (**A**) Protomers were aligned to residues 25-35 in protomer A to superimpose the core of the aptamer as there is some variation in the P1 helix. The all atom RMSD. comparing protomers B, C, and D to protomer A ranges from 4.8 to 7.1 Å. This large RMSD. between protomers is due to conformational flexibility of P1 relative to the core of the RNA. (**B**) The P1 helix was removed from each protomer as this region varied greatly among the protomers. An all atom RMSD. of the core of protomers B, C, and D with protomer A (residues 6-41) ranged from 0.97 to 1.3 Å indicating good agreement.

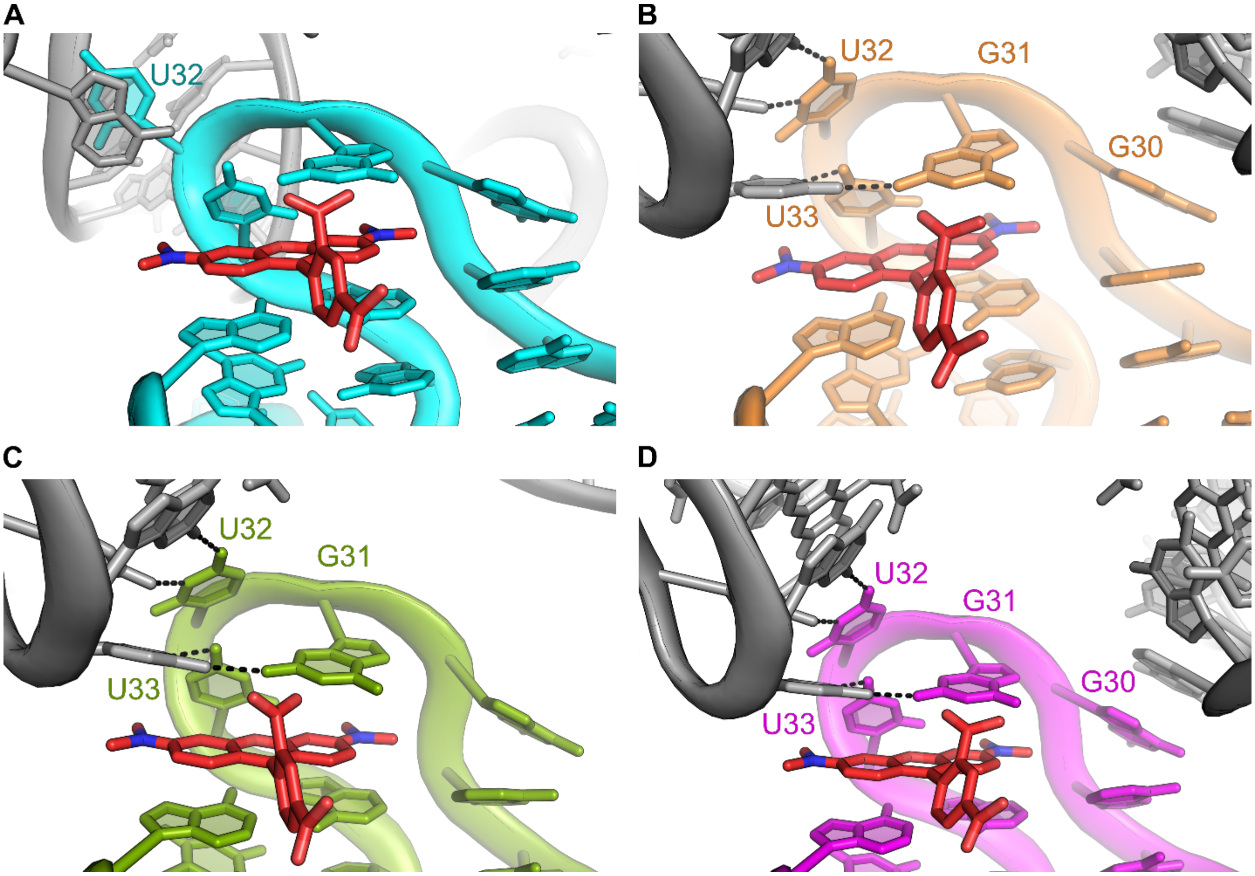

**Supplemental Figure S5.** Lattice interactions at binding site for all protomers in RhoBAST:5-TAMRA structure. Neighboring RNA molecules are shown in gray. (**A**) Protomer A binding site. The lattice interactions of protomer A (cyan) consists of U32 π-stacking with an adjacent adenine base. No other interactions occur with important binding site residues, so this protomer was selected for analysis. (**B**) Protomer B binding site. Protomer B (orange) has lattice contact interactions with G30, G31, U32, and U33 of the binding pocket. (**C**) Protomer C binding site. Protomer C (green) has lattice contact interactions with G31, U32, and U33 of the binding pocket. (**D**) Protomer D binding site. Protomer D (magenta) has lattice contact interactions with G30, G31, U32, and U33 of the binding site.

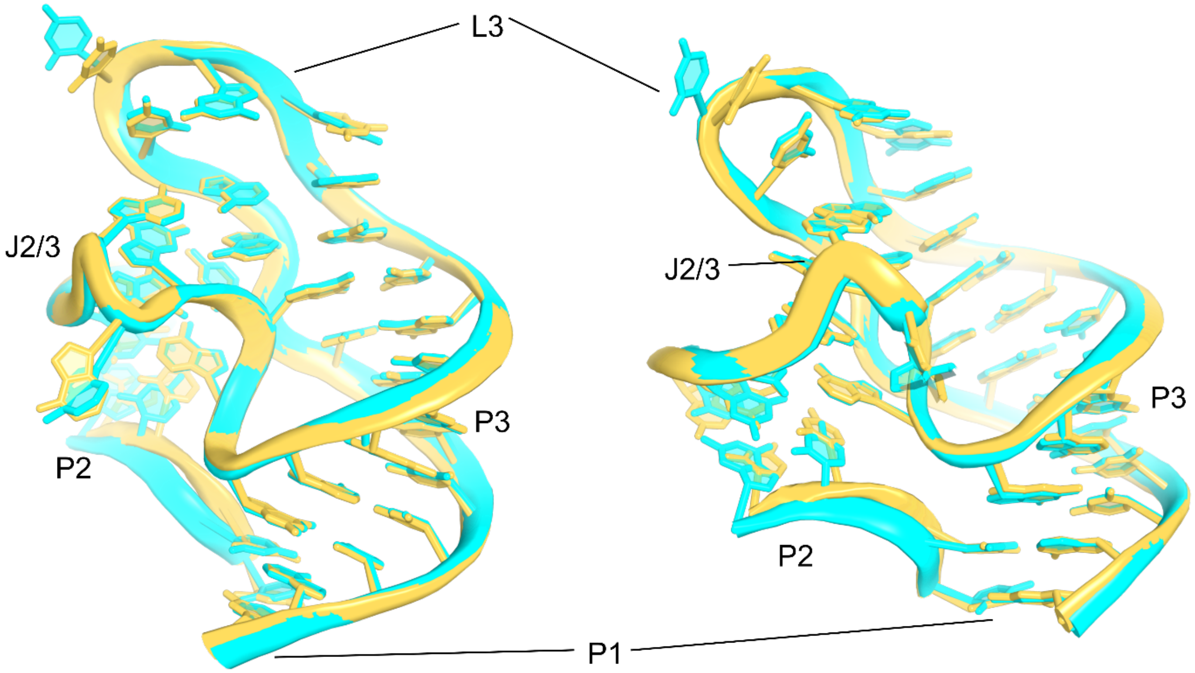

**Supplemental Figure S6.** Global alignment of RhoBAST:5-TAMRA structure with RhoBAST:TMR-DN. An all-atom superimposition was performed using the RhoBAST:5-TAMRA structure shown in cyan (PDB ID: 9BUN and chain A using residues 4-7 and 12-44) and the RhoBAST:TMR-DN structure (3) shown in yellow (PDB ID: 8JY0 and chain B using residues 6-9 and 28-60). Because the sequences of helices P1 and P2 differed between constructs, these variable sequences were removed to compare the core RNA structure that is identical in both constructs. The calculated all atom RMSD. of the core is 1.07 Å.

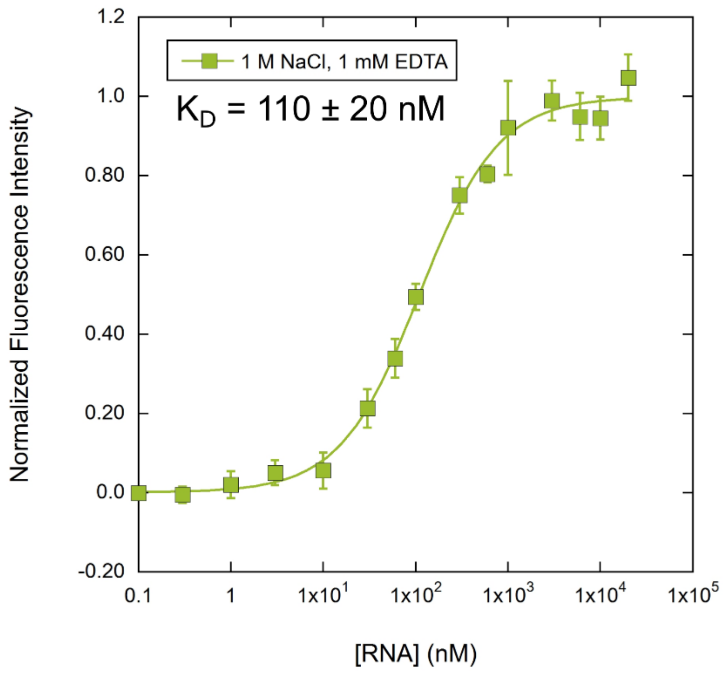

**Supplemental Figure S7.** Fluorescence binding assay of RhoBAST binding to SpyRho555 in high monovalent cation concentrations without $Mg^{2+}$. Binding assays were performed as described in Methods except the final buffer concentration was 20 mM HEPES pH 7.4, 1 M NaCl, 1 mM EDTA, and 0.01% v/v Tween-20. Data are shown as mean and s.d. from independent measurements (n = 3).

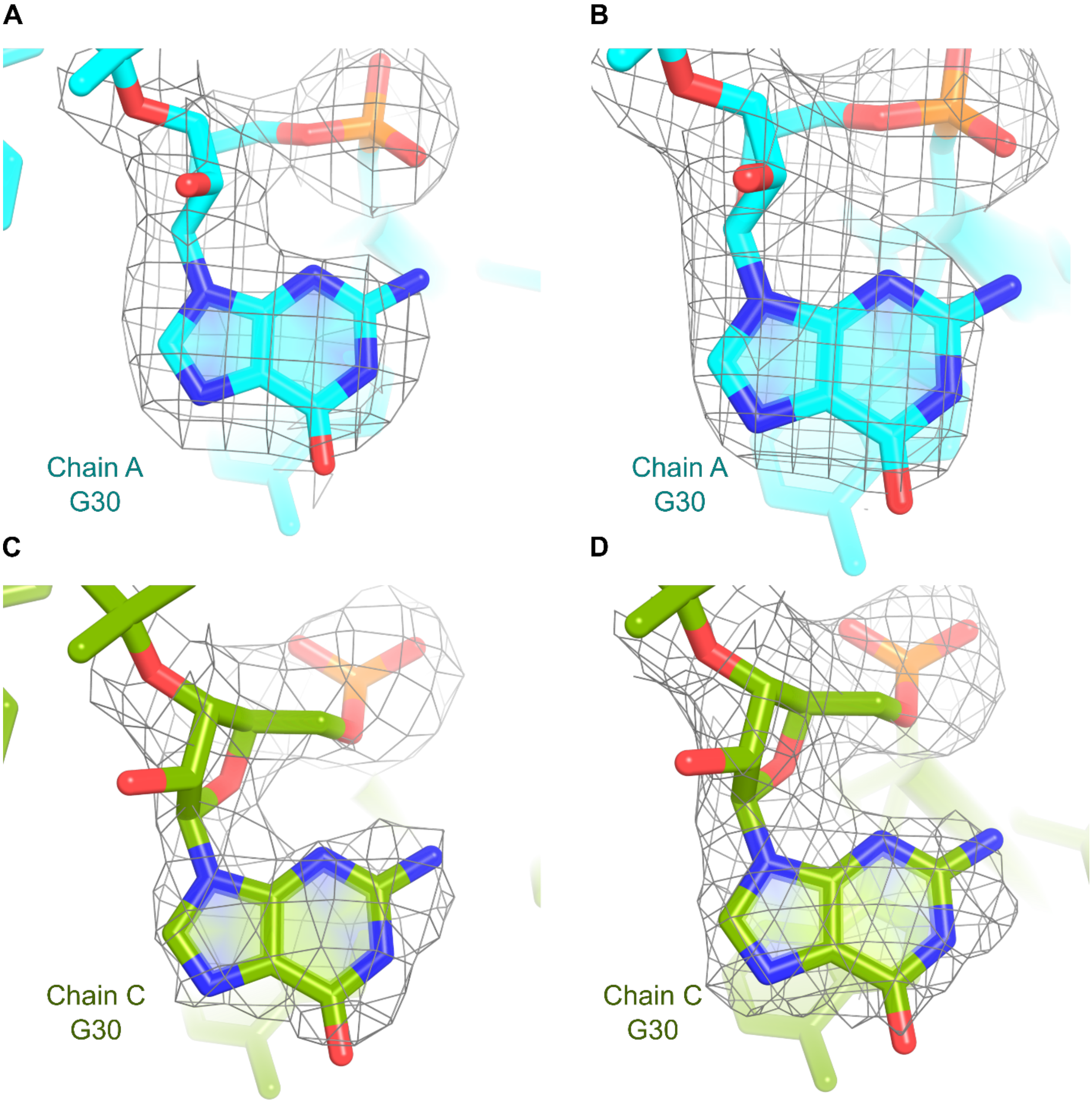

**Supplemental Figure S8.** *Syn* configuration of G30 in RhoBAST:5-TAMRA structure. The G30 nucleotide does not contain π-stacking interactions from lattice contacts in chain A (cyan) and chain C (green). The assignment of the *syn* configuration was examined by looking at the density modified map obtained just after phasing (1) or a simulated-annealing composite omit map (2). The $2|F_o| - |F_c|$ density is carved to 2 Å around residue 30 at 1 σ. (**A**) G30 shown in chain A with the density modified map. (**B**) G30 shown in chain A with the composite omit map. (**C**) G30 shown in chain C with the density modified map. (**D**) G30 shown in chain C with the composite omit map.

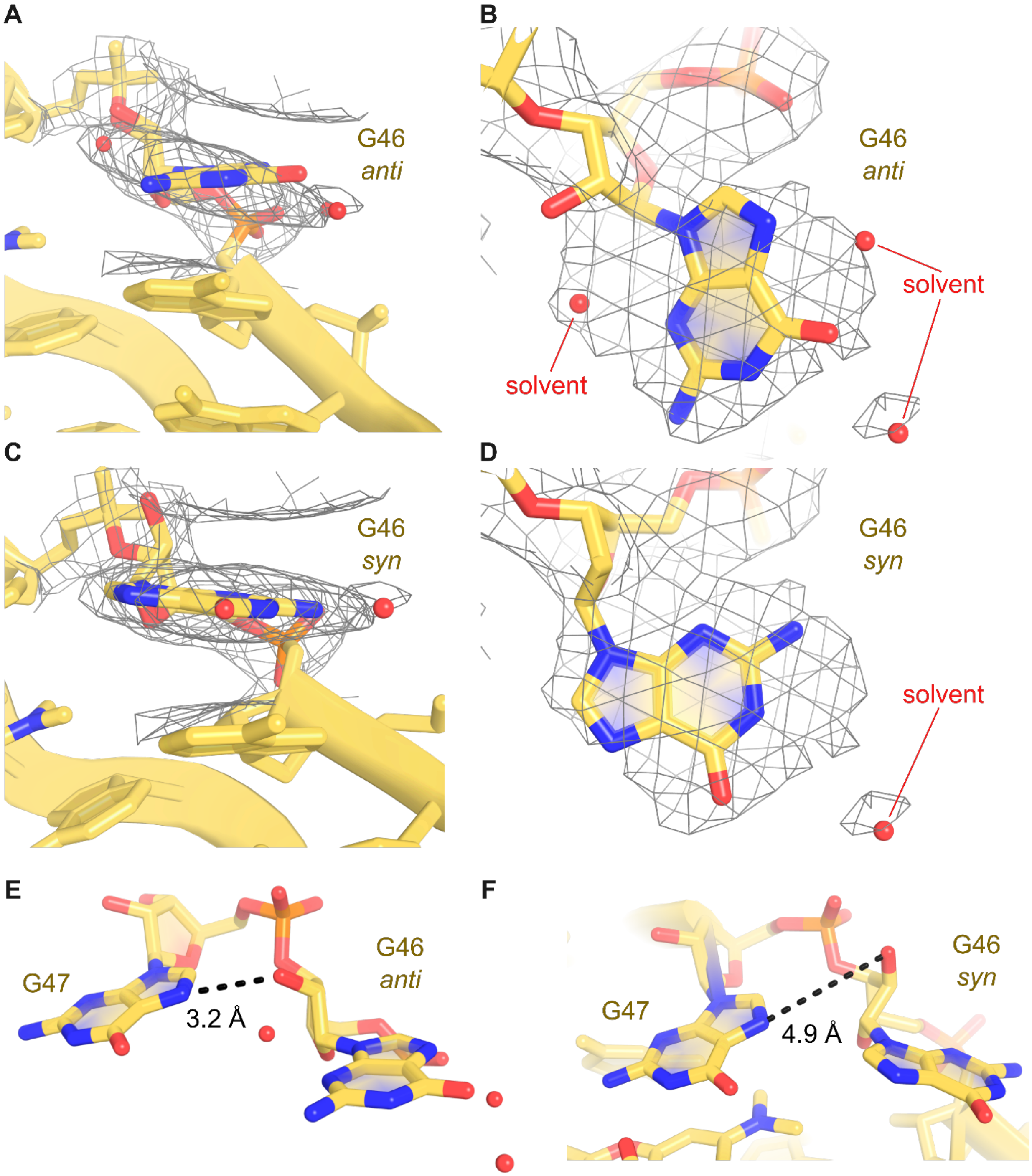

**Supplemental Figure S9.** G30 configuration in RhoBAST:TMR-DN structure. The equivalent G30 position in the RhoBAST:TMR-DN model (PDB ID: 8JY0) (3) is G46. The 2|$F_o$| − |$F_c$| density was obtained from the PDB and carved 3 Å around residue 46 at 1 σ. Protomer B, the protomer discussed by the authors of the RhoBAST:TMR-DN structure (3), is shown here colored in yellow. Solvent molecules are shown as nonbonded spheres within 5 Å of G46. (**A**) The *anti* configuration of G46 as depicted in protomer B from the side. (**B**) The *anti* configuration of G46 as depicted in protomer B from the top. (**C**) The *syn* configuration of G46 in protomer B was modelled by removing two nearby solvent molecules (301 and 302), manually adjusting the G46 nucleotide, and performing a Real Space Refine Zone in Coot (4) against the 2|$F_o$| − |$F_c$| density. The side view of the *syn* configuration demonstrates much better agreement with the electron density. (**D**) Top view of the *syn* configuration of protomer B displays good agreement with the electron density. (**E**) The authors reported a hydrogen bond with a distance of 3.2 Å occurring between the 2′-hydroxyl of G46 (*anti*) and the N7 of G47 in the RhoBAST:TMR-DN structure (3). (**F**) In protomer B, the *syn* configuration precludes the interaction between G46 and G47 with a distance of 4.9 Å.

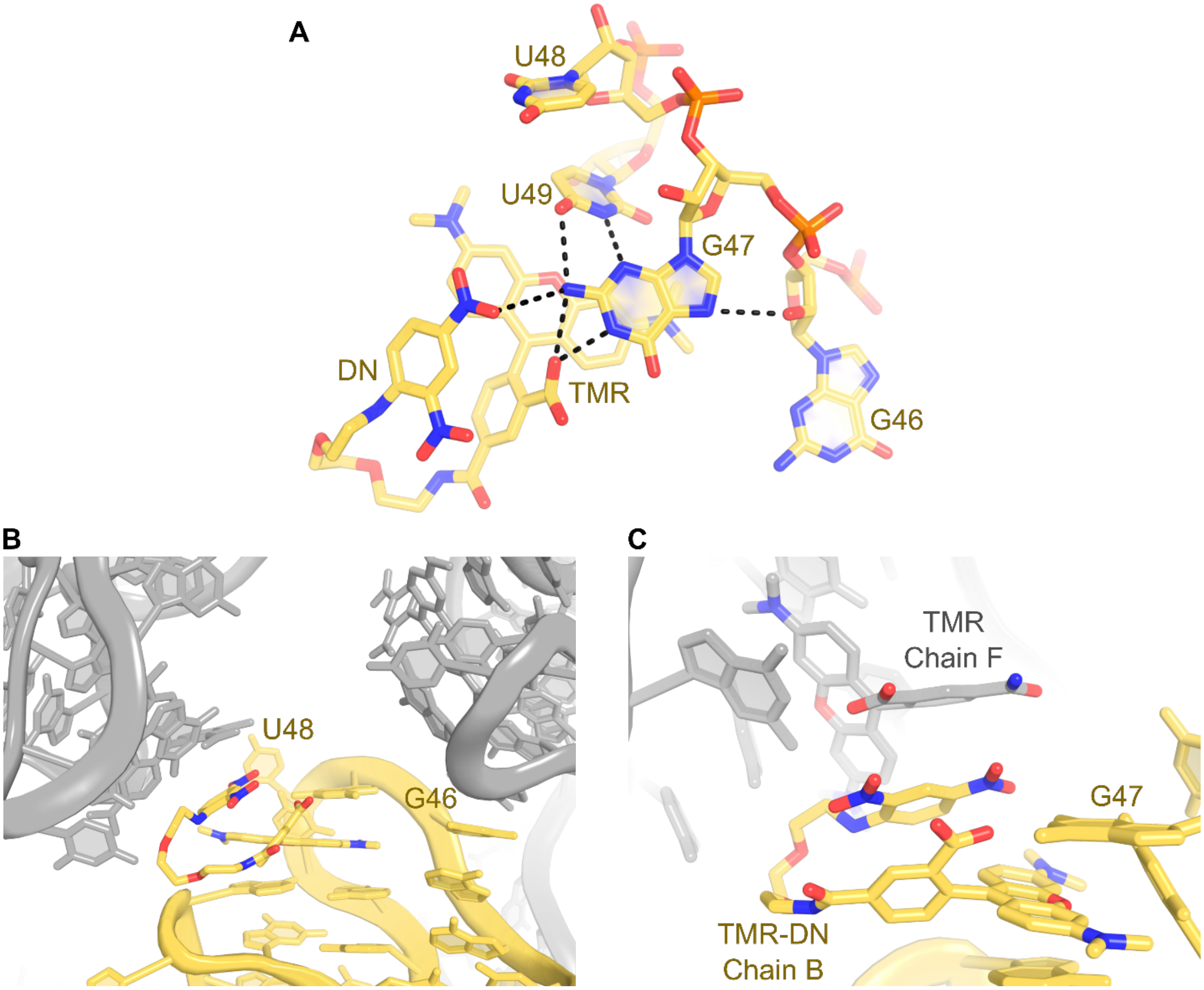

**Supplemental Figure S10.** Ligand interactions in the RhoBAST:TMR-DN structure (PDB ID: 8JY0) and lattice interactions. (**A**) Ligand interactions as described by the authors of the RhoBAST:TMR-DN structure (3). The ligand, TMR-DN, interacts with the WCF face of G47 through its 3-carboxylate of TMR and nitro group of DN. The WCF edge of U49 interacts with the sugar edge of G47. The 2′-hydroxyl of G46 in the *anti* configuration interacts with N7 of G47. (**B)** Overview of lattice interactions in protomer B near the binding site in the RhoBAST:TMR-DN structure. A neighboring RNA molecule (gray) stacks with G46 of protomer B (yellow) while the binding site and ligand of another protomer stacks with the TMR-DN ligand of protomer B. (**C**) Close-up view of lattice interactions with TMR-DN ligand of protomer B. The DN moiety of the TMR-DN ligand in protomer B directly stacks with the phenyl ring of the TMR ligand in protomer F. These lattice interactions influence the binding pocket of protomer B discussed in the RhoBAST:TMR-DN paper (3).

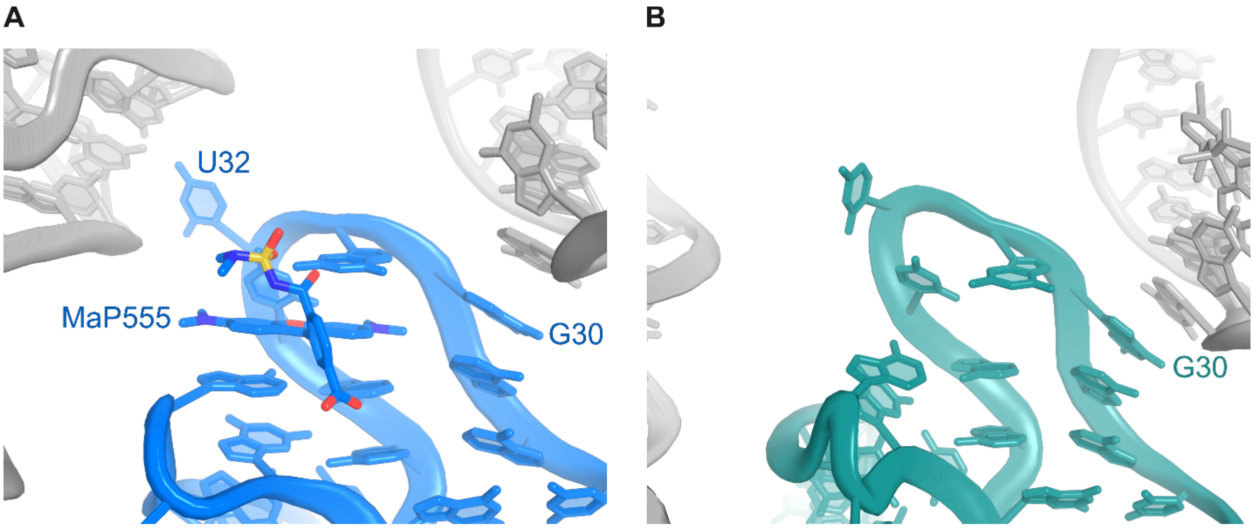

**Supplemental Figure S11.** Lattice interactions near binding site in RhoBAST:MaP555 structure. (**A**) Lattice interactions of protomer A in the RhoBAST:MaP555 structure (PDB ID: 9DXL) shown as cartoons with the ligand as sticks. G30 and U32 of protomer of A (blue) interact with neighboring RNA molecules (gray). (**B**) Lattice interactions of protomer B in the RhoBAST:MaP555 structure shown as cartoons. G30 of protomer B (teal) stacks with a neighboring RNA molecule (gray) adjacent to the binding site. The electron density in the binding pocket of protomer B is poor, so the ligand was not placed in this protomer. The data quality cannot unambiguously assign protomer B as the apo configuration.

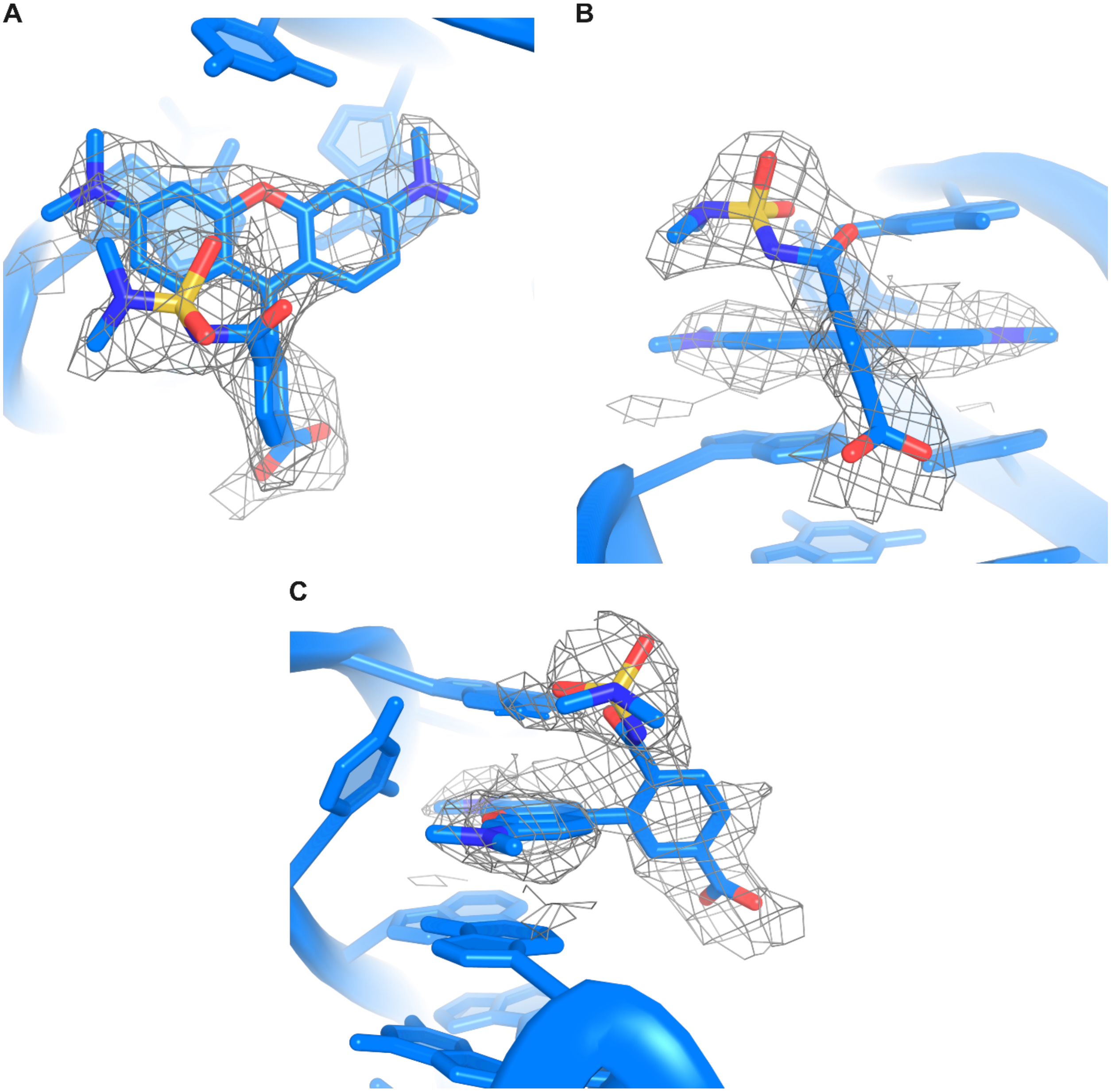

**Supplemental Figure S12.** Electron density supporting placement of MaP555 in RhoBAST:MaP555 structure (PDB ID: 9DXL). A simulated-annealing composite omit map (2) was generated for protomer A bound to MaP555. The 2|$F_o$| − |$F_c$| composite omit map is carved 2 Å around MaP555 and shown at 1 σ. (**A**) Front view of MaP555 electron density. (**B**) Top-down view of MaP555 electron density. The G31 base has been removed for clarity. (**C**) Side view of MaP555 electron density.

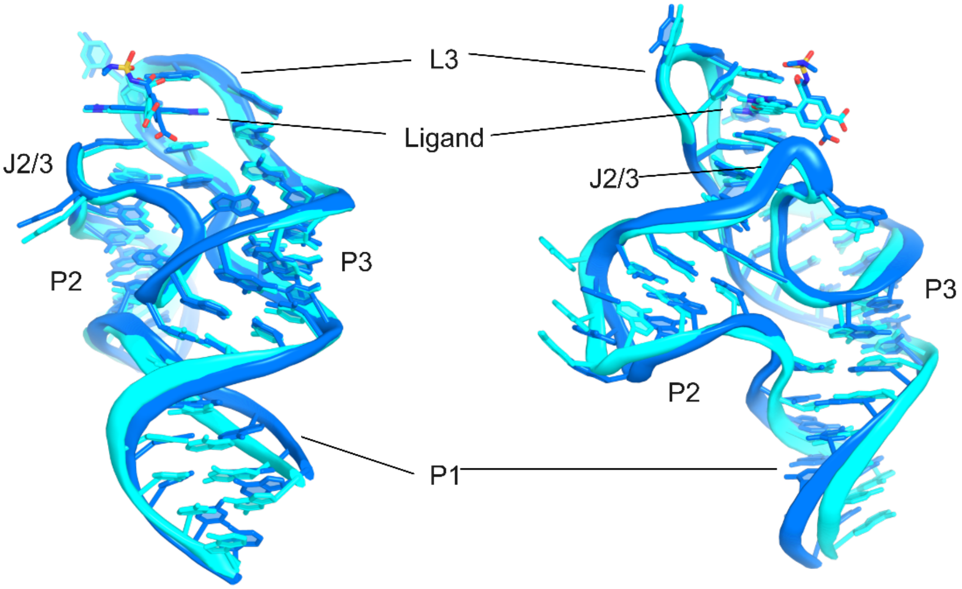

**Supplemental Figure S13.** Superimposition of RhoBAST:5-TAMRA and RhoBAST:MaP555 structures. Protomer A of the RhoBAST:5-TAMRA structure (cyan, PDB ID: 9BUN) was superimposed with protomer A of the RhoBAST:MaP555 structure (blue, PDB ID: 9DXL). Both protomers demonstrate good agreement with each other and have an all atom RMSD of 1.9 Å.

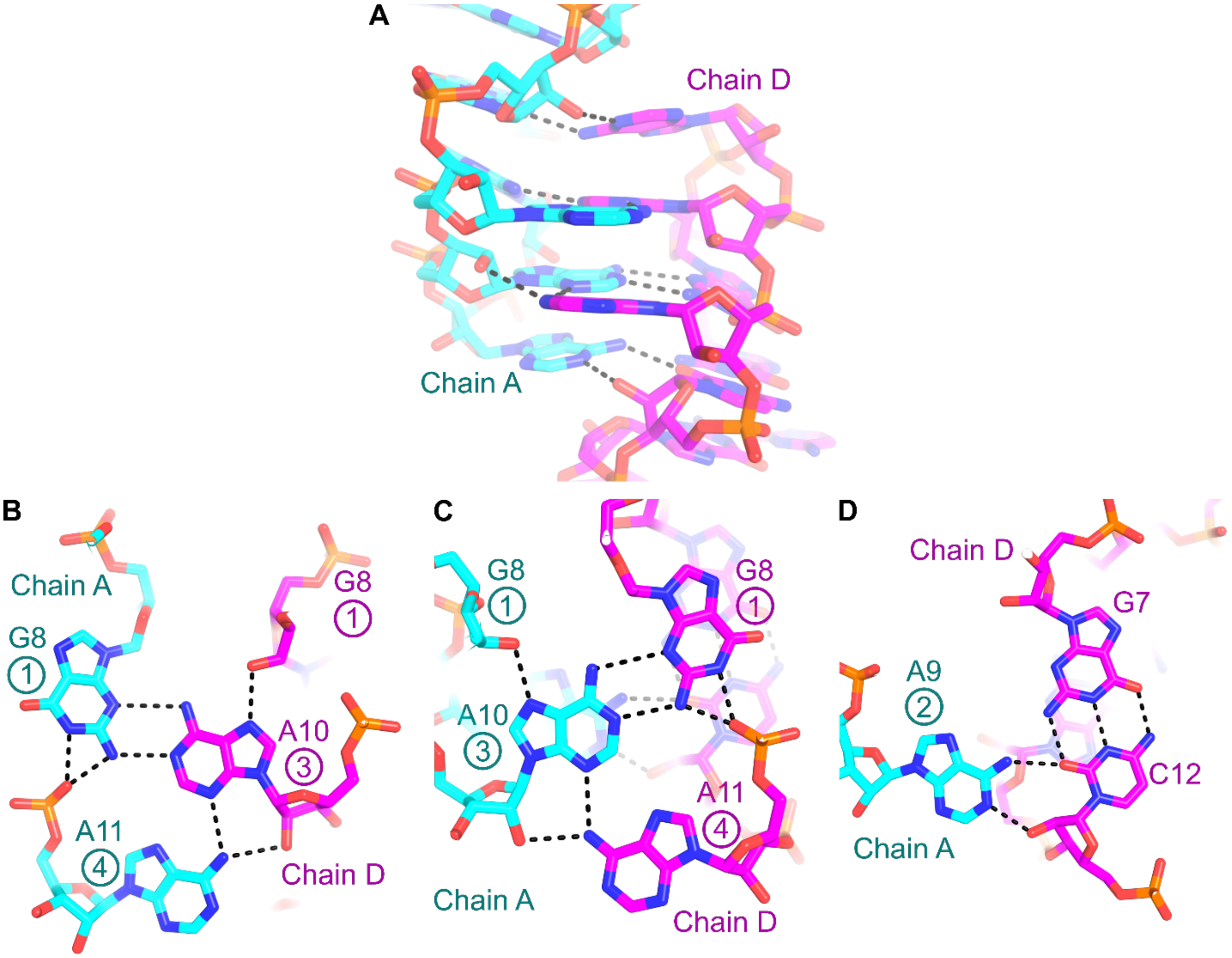

**Supplemental Figure S14.** GAAA tetraloop interactions between chain A and chain D in RhoBAST:5-TAMRA structure. (**A**) Overall interactions between terminal P2 loops of chain A (cyan) and chain D (magenta). Extensive interactions between these two protomers can be found in the crystal lattice. (**B**) The first (G8) and fourth (A11) residues in the GNRA tetraloop of chain A no longer interact directly with each other and form a base triple with the third residue of the GNRA tetraloop in chain D. The position of the residue within the tetraloop is denoted by the circled number. (**C**) The third residue of the tetraloop (A10) in chain A forms a base triple with residues 1 and 4 of the chain D tetraloop. (**D**) The second residue of the chain A tetraloop (A9) interacts with the distal WCF base pair of P2 (G7-C12) in chain D through its WCF edge and contacts the sugar edge and 2′-hydroxyl of C12 in chain D.

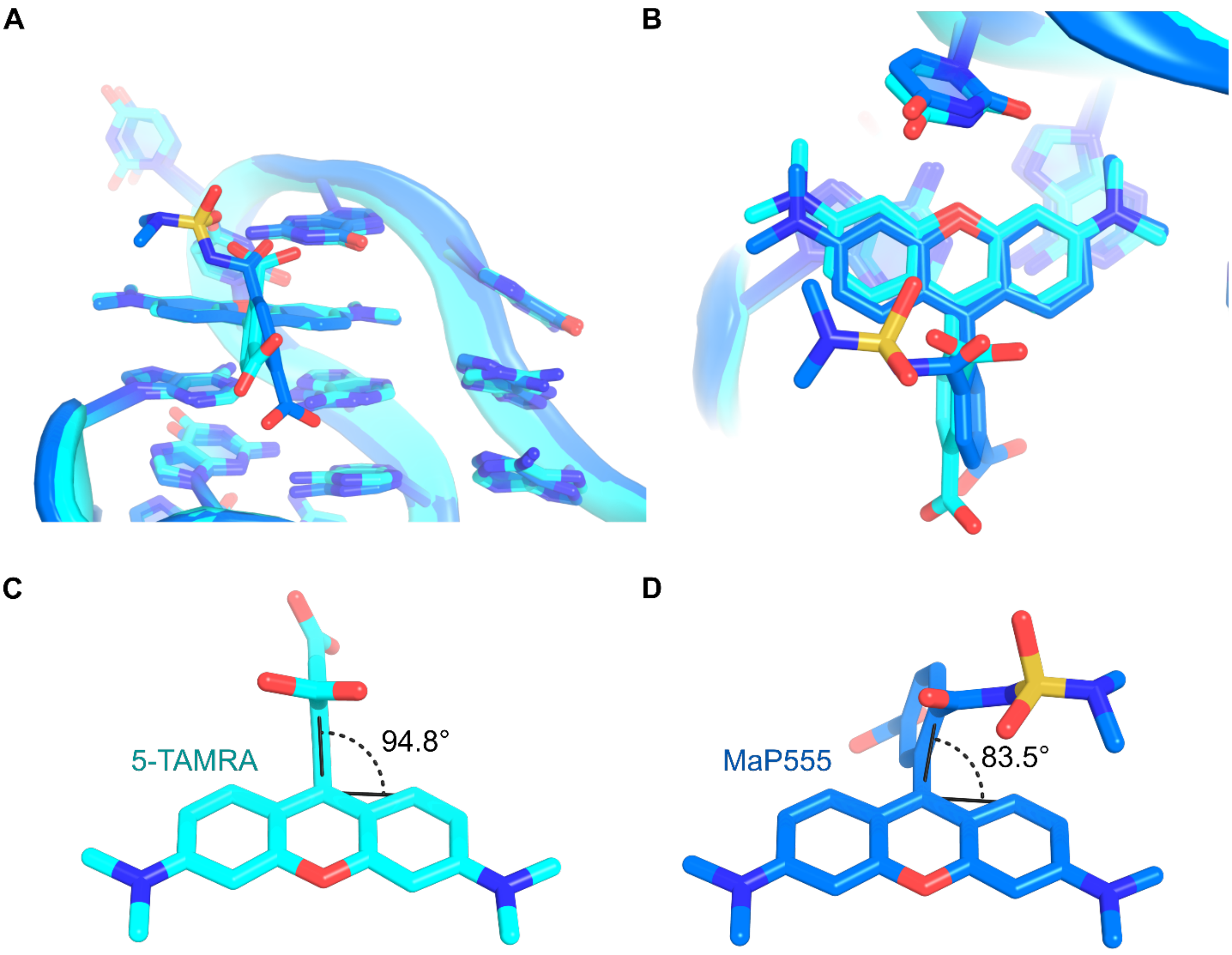

**Supplemental Figure S15.** Comparison of 5-TAMRA and MaP555 bound to RhoBAST. The RhoBAST:5-TAMRA structure is shown in cyan while the RhoBAST:MaP555 structure is shown in blue. (**A**) Superimposition of the ligand binding pocket. 5-TAMRA and MaP555 are bound in the same location. (**B**) Top-down view of both ligands. The xanthene core of 5-TAMRA and MaP55 are positioned almost identically between structures although MaP555 is slightly rotated away from the RNA. The buried surface area (BSA) for 5-TAMRA was calculated as 336 Å$^2$ compared to MaP555 which has a BSA of 327 Å$^2$. The G31 base has been removed for clarity. (**C**) Rotation of phenyl moiety in 5-TAMRA. The angle between the 3-1-1′ positions of 5-TAMRA was measured to determine the degree of rotation relative to the planar xanthene group. This measurement for 5-TAMRA is 94.8°. (**D**) Rotation of phenyl moiety in MaP555. The same measurement taken between the 3-1-1′ positions of MaP555 was determined as 83.5°. The difference between these angles for 5-TAMRA and MaP555 is 11.3°.

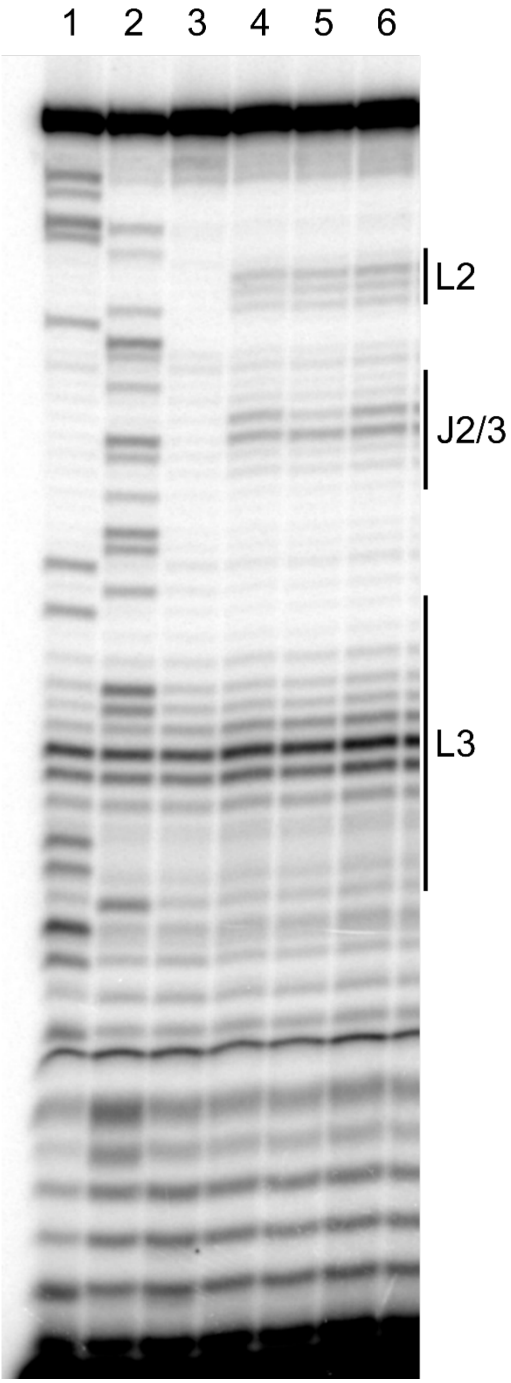

1. C sequencing
2. G sequencing
3. DMSO
4. NMIA
5. 10 µM 5(6)-TAMRA
6. 10 µM SpyRho555

**Supplemental Figure S16.** Full SHAPE chemical probing sequencing gel of WT RhoBAST.

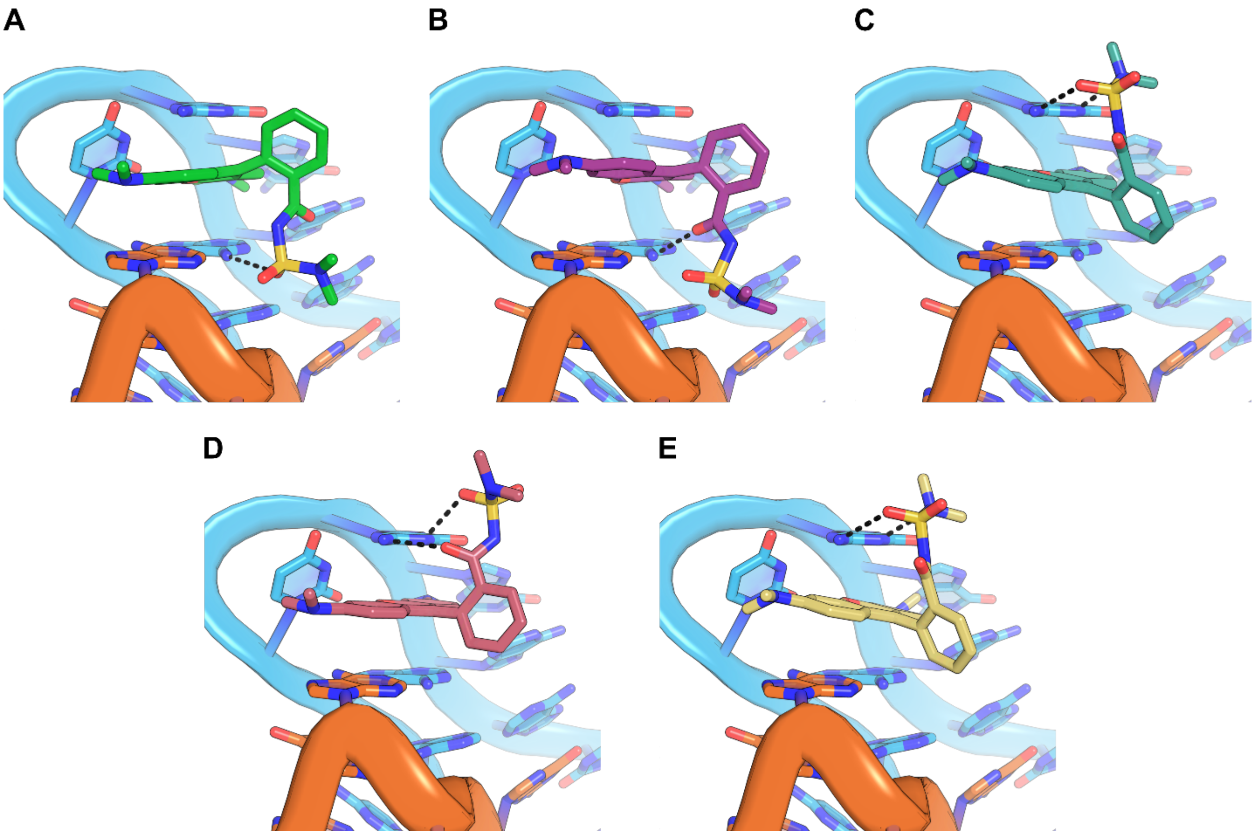

**Supplemental Figure S17.** Docking analysis of the open configuration of SpyRho555. The top five ranked docking posed using the open configuration of SpyRho555 are displayed with hydrogen bonding interactions. (**A**) Top-ranked pose of the open form of SpyRho555. (**B**) Second-ranked pose of the open form of SpyRho555. (**C**) Third-ranked pose of the open form of SpyRho555. (**D**) Fourth-ranked pose of the open form of SpyRho555. (**E**) Fifth-ranked pose of the open form of SpyRho555.

**A** **B** **C**

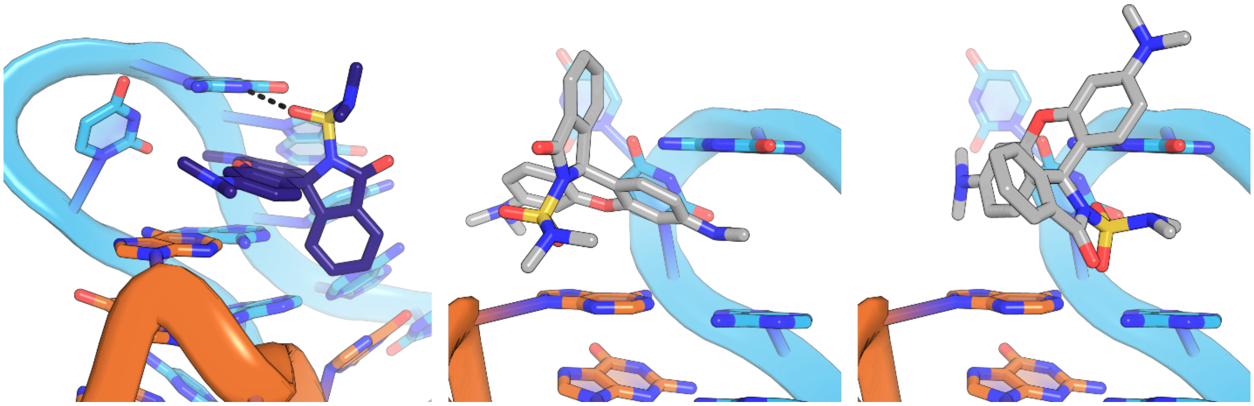

**Supplemental Figure S18.** Docking analysis of the closed configuration of SpyRho555. (**A**) Third-ranked pose of the closed form of SpyRho555 with hydrogen bonding interactions shown. (**B**) Fourth-ranked pose of the closed form of SpyRho555. The xanthene ring is pushed out of the binding pocket inconsistent with the crystal data and excluded from discussions. (**C**) Fifth-ranked pose of the closed form of SpyRho555. The 3-position group of SpyRho555 is located between G31 and the purine floor while the xanthene does not stack with the RNA. This docking result was excluded from discussions.

**Supplemental Table S1.** Sequences used in this study. The T7 promoter is highlighted in yellow, mutations to the WT sequences are highlighted in cyan, and the SHAPE cassette is highlighted in green.

| Name | Description | Sequence (5′ to 3′) |
|---|---|---|
| 5′ GEN | 5′ primer for all PCRs | GCGCGCGAATTCTAATACGACTCACTATAG |
| RhoBAST WT | WT RhoBAST construct | GCGCGCGAATTCTAATACGACTCACTATAGGAACCTCCGCGAAAGCGGTGAAGGAGAGGCGCAAGGTTAACCGCCTCAGGTTCC |
| 3′ RhoBAST | General 3′ primer for RhoBAST constructs | GGAACCTGAGGCGGTTAACCTTG |
| RBG31A | RhoBAST G31A mutant | GGCTAATACGACTCACTATAGGAACCTCCGCGAAAGCGGTGAAGGAGAGGCGCAAGATTAACCGCCTCAGGTTCC |
| 3′ RBG31A | 3′ primer for G31A mutant | GGAACCTGAGGCGGTTAATCTTG |
| RBG31C | RhoBAST G31C mutant | GGCTAATACGACTCACTATAGGAACCTCCGCGAAAGCGGTGAAGGAGAGGCGCAAGCTTAACCGCCTCAGGTTCC |
| 3′ RBG31C | 3′ primer for G31C mutant | GGAACCTGAGGCGGTTAAGCTTG |
| RBG19C | RhoBAST G19C mutant | GGCTAATACGACTCACTATAGGAACCTCCGCGAAAGCGGTGAAGCAGAGGCGCAAGGTTAACCGCCTCAGGTTCC |
| RBG19C/C27G | RhoBAST G19C & C27G mutant | GGCTAATACGACTCACTATAGGAACCTCCGCGAAAGCGGTGAAGCAGAGGCGGAAGGTTAACCGCCTCAGGTTCC |
| 3′ RBC27G | 3′ primer for C27G mutant | GGAACCTGAGGCGGTTAACCTTC |
| RBU14C/A20G | RhoBAST U14C & A20G mutant | GGCTAATACGACTCACTATAGGAACCTCCGCGAAAGCGGCGAAGGGGAGGCGCAAGGTTAACCGCCTCAGGTTCC |
| RBU14G/A20C | RhoBAST U14G & A20C mutant | GGCTAATACGACTCACTATAGGAACCTCCGCGAAAGCGGGGAAGGCGAGGCGCAAGGTTAACCGCCTCAGGTTCC |
| RBC4U/G44A | RhoBAST C4U & G44A mutant | GCGCGCGAATTCTAATACGACTCACTATAGGAACTTCCGCGAAAGCGGTGAAGGAGAGGCGCAAGGTTAACCGCCTCAAGTTCC |
| RBC6A/G13U | RhoBAST C6A & G13U mutant | GCGCGCGAATTCTAATACGACTCACTATAGGAACCTACGCGAAAGCGTTGAAGGAGAGGCGCAAGGTTAACCGCCTCAGGTTCC |
| RBP1-4/P2-2 | RhoBAST construct for crystallization | GGCTAATACGACTCACTATAGGACTCGGAAACGTGAAGGAGAGGCGCAAGGTTAACCGCCTCAGTCCA |
| 3′ RBP1-4/P2-2 | 3′ primer for RhoBAST crystal construct | TGGACTGAGGCGGTTAACCTTGC |
| 3′ RBP1-4/P2-2 2′-*O*-methyl | 3′ primer for RhoBAST crystal construct containing 2′-*O*-methyl | mUmGGACTGAGGCGGTTAACCTTGC |

| | | |
|---|---|---|
| | modifications, used for crystallization | |
| 3′ RB + SHAPE cassette | 3′ primer for SHAPE chemical probing of RhoBAST | GAACCGGACCGAAGCCCGGGAACCTGAGGCGGTTAACCTTG |

**Supplemental Table S2.** Data collection and model statistics.

| Crystal | *RhoBAST:5-TAMRA* | *RhoBAST:MaP555* |
|---|---|---|
| PDB ID | 9BUN | 9DXL |
| **Data collection** | | |
| Wavelength (Å) | 1.09720 | 1.54178 |
| Resolution Range (Å)[a] | 44.45-1.95 (2.00-1.95) | 20.00-2.80 (2.90-2.80) |
| Space Group | C 2 2 $2_1$ | P 1 $2_1$ 1 |
| Unit Cell | | |
| a, b, c (Å) | 102.72, 175.21, 88.69 | 38.54, 54.86, 83.50 |
| α, β, γ (°) | 90.00, 90.00, 90.00 | 90.00, 103.37, 90.00 |
| Total Reflections | 3750599 (188221) | 318867 (30360) |
| Unique Reflections | 58528 (4067) | 8812 (839) |
| $R_{meas}$ | 0.090 (8.201) | 0.127 (0.352) |
| $R_{pim}$ | 0.013 (1.180) | 0.074 (0.210) |
| Multiplicity | 64.1 (46.3) | 2.7 (2.4) |
| Completeness (%) | 99.9 (99.7) | 98.0 (96.2) |
| I/σ(I) | 34.2 (0.6) | 10.0 (2.7) |
| $CC_{1/2}$ | 1.00 (0.319) | 0.948 (0.815) |
| **Refinement** | | |
| Resolution (Å) | 44.245 – 2.100 (2.175– 2.100) | 19.27-2.80 (2.97-2.80) |
| No. reflections | 46132 (4521) | 8303 (800) |
| $R_{work}/R_{free}$ | 0.216 (0.316)/0.241 (0.357) | 0.254 (0.353) / 0.271 (0.409) |
| No. non-hydrogen atoms | 4840 | 2153 |
| RNA | 4144 | 3118 |
| Ligand | 212 | 65 |
| Ions | 375 | 9 |
| Water | 463 | 34 |
| B-factors | | |
| RNA | 64.83 | 79.32 |
| Ligand | 52.21 | 45.09 |
| Ions | 91.95 | 58.52 |
| Water | 61.75 | 39.01 |
| r.m.s. deviations | | |
| Bond Length (Å) | 0.003 | 0.001 |
| Bond Angle (°) | 0.627 | 0.495 |

[a]Values in parentheses are for the highest resolution shell.

**Supplemental Table S3.** Thermodynamic values for wild type RhoBAST binding to 5-TAMRA determined using isothermal titration calorimetry.

| **$K_D$ ± s.d. (nM)** | **N value ± s.d.** | **ΔH ± s.d. (kcal/mol)** | **-TΔS ± s.d. (kcal/mol)** |
|---|---|---|---|
| 70 ± 20 | 0.80 ± 0.05 | -21 ± 1 | 12 ± 1 |

**Supplemental Table S4.** Binding pocket analysis of RhoBAST with top docking poses.

| Ligand | RNA Model | Description | 3-postion group of ligand pointing toward G31? | Buried Surface Area (Å$^2$) |
|---|---|---|---|---|
| 5-TAMRA | RhoBAST:5-TAMRA structure | Crystal structure | Yes | 336 |
| MaP555 | RhoBAST:MaP555 structure | Crystal structure | Yes | 327 |
| Open SpyRho555 | RhoBAST:MaP555 | Crystal structure with 6-carboxylate of MaP555 removed | Yes | 328 |
| Open SpyRho555 | RhoBAST:5-TAMRA structure | Docking pose 1 | No | 409 |
| Open SpyRho555 | RhoBAST:5-TAMRA structure | Docking pose 2 | No | 412 |
| Open SpyRho555 | RhoBAST:5-TAMRA structure | Docking pose 3 | Yes | 348 |
| Open SpyRho555 | RhoBAST:5-TAMRA structure | Docking pose 4 | Yes | 348 |
| Open SpyRho555 | RhoBAST:5-TAMRA structure | Docking pose 5 | Yes | 349 |
| Closed SpyRho555 | RhoBAST:5-TAMRA structure | Docking pose 1 | Yes | 339 |
| Closed SpyRho555 | RhoBAST:5-TAMRA structure | Docking pose 2 | No | 376 |
| Closed SpyRho555 | RhoBAST:5-TAMRA structure | Docking pose 3 | Yes | 348 |
